# Ancestral and environmental diversity shape the immune landscape in Indonesia

**DOI:** 10.64898/2026.02.15.704933

**Authors:** Muhamad Fachrul, Pongsakorn Sukonthamarn, Pradiptajati Kusuma, Monika Meili Novita, Isabela Alvim, Isabella Apriyana, Chelzie Crenna-Darusallam, Andreas Christian, Alice Groudko, Robert Kendle, Prisca Cynthia Limardi, Evan D Mee, Sukma Oktavianthi, Lance M Peter, Lidwina Priliani, Bertha Letizia Utami, Fredrik Sokoy, Simon Abdi K. Frank, Desak Made Wihandani, Ni Nyoman Ayu Dewi, Agus Eka Darwinata, Murray P. Cox, Nicholas E. Banovich, Herawati Sudoyo, Safarina G. Malik, Irene Gallego Romero

## Abstract

Island Southeast Asia (ISEA) remains consistently underrepresented in human genomic resources despite its exceptional ancestral and lifestyle diversity. The interplay between the region’s complex population history and its environmental variation provide a window into how ancestry and environment jointly shape the human immune system. Here we report the generation of single-cell PBMC profiles from 199 Indonesians sampled across four communities in the islands of Bali and New Guinea. These groups capture diversity in regional genetic ancestries (East Asian-like and Papuan-like) and lifestyle contrasts (urban versus rural communities in Bali; highland versus lowland communities in New Guinea). We identify over 4,000 expression quantitative trait loci (eQTLs) across nine immune cell types, including eQTLs driven by introgression from both Neanderthals and Denisovans at genes such as *IL7R*, *HLA-E*, or *STAT2*. We also find evidence of local ancestry driving gene-by-environment interactions at pathogen receptors such as *MARCO*, although the majority of gene-by-environment interactions are not driven by differences in genetic structure between populations. Beyond direct genetic effects, we construct gene co-expression networks that consistently identify environmental signatures, as well as T-cell receptor repertoires that distinguish specific communities, with excess representation of interferon-stimulated genes in rural, but not urban samples. This work establishes a framework for population-aware functional genomics in understudied regions and highlights how ancestral and environmental diversity jointly shape human immunity in this globally important yet underrepresented region.

## Introduction

Indonesia is the world’s fourth largest country by population, and home to approximately 280 million people. Spanning over 17,000 islands in Island Southeast Asia (ISEA), the country and the region are host to significant levels of cultural, linguistic and genetic diversity [1, 2], but they both remain significantly underrepresented in genomic resources, even relative to other non-European-like genetic ancestries. The first Indonesian whole genome sequences were only published in 2019 [3, 4] and functional genomics data, especially from healthy donors, remains equally sparse, with the largest dataset including 117 individuals sampled across three islands [5]. Given this scarcity, studies linking genotype and phenotype through formal identification of expression quantitative trait loci or similar also remain rare [6], despite their potential to inform on the importance of gene-by-environment interactions. This scarcity limits our collective ability to interpret how population history, local ancestry, and environmental diversity shape cell type-specific regulation. Indonesia’s lower-lying and coastal regions have a high pathogen load and diseases like tuberculosis [7], malaria [8], and dengue [9] remain endemic, while life in the interior can necessitate adaptation to high altitudes, with parts of Java and New Guinea Island being over 2000 meters above sea level [10, 11]. The region’s lengthy and complex history of human habitation, which dates back to its initial settlement and includes deep and region-specific genetic structure [12, 13, 14, 15] and multiple archaic introgression events not present anywhere else in the world [16, 3], means that today Indonesia is characterised by a longitudinal gradient of genetic admixture, whereby individuals from the eastern part of the country are exclusively or primarily of Papuan-like genetic ancestry, which transitions into East Asian-like genetic ancestry moving westward [2]. The combination of genetic and environmental diversity that characterises the region makes ISEA a compelling model for investigating the interplay of genetic and environmental forces in shaping human immune diversity.

When individuals from non-European-like genetic ancestries are included in studies they are frequently sampled in urbanised settings, or as members of diaspora or migrant communities living in Europe or North America, which provides limited representation of genetic and environmental diversity [17, 18]. These gaps have consequences for understanding immunity and diseases in tropical and high-pathogen settings, such as Indonesia and ISEA. The recently published Asian Immune Diversity Atlas (AIDA, [19]), provided, for the first time, expression data at a single-cell resolution across multiple populations and sites spanning Southeast, South and East Asia, including 59 Thai donors living in Bangkok, the capital of Thailand, and 206 donors from Singapore, a tropical high income country (HIC). This work identified significant differences in immune cell type proportions across sampling sites and ancestries, including differences across mainland Southeast Asian populations, but did not include any ISEA populations. A separate study that focused on immune cell repertoire diversity across individuals in the Netherlands and Senegal living in urban and rural settings recently reported differences in immune cell subpopulations and metabolic states across environments, and was able to replicate these observations in a cohort of 8 Indonesians living in rural and urban settings [20], again pointing towards an impact of the environment on immune cell state.

Single-cell resolution immune atlases that integrate diversity therefore provide a strong way to map how ancestry, environment, and cellular gene expressions interact with immune phenotypes. Here, we present a single-cell resolution atlas of gene expression in whole blood from 199 individuals across four sites in two Indonesian islands, Bali and Papua, constituting the first single-cell resolution immune atlas from ISEA. This dataset reflects Indonesian and ISEA priorities and realities that are absent from the vast majority of existing public genomic resources, including representation of both regional genetic ancestries, and spanning a lifestyle gradient from rural subsistence agriculture to a sizeable Global South city. This allows us to examine how deep population history, recent demographic processes, and environment in these underrepresented but diverse populations jointly shape the immune phenotypes. Our work demonstrates both the feasibility of conducting functional genomics research in resource-limited settings as well as its attendant challenges, and provides a road-map for population-aware functional genomics in underrepresented populations worldwide.

## Results

### Capturing ancestral and environmental diversity in Indonesia at single-cell resolution

To atlas circulating immune cell diversity across Indonesia we collected whole blood samples from 199 donors across four sites in the Indonesian archipelago and generated single-cell RNA-sequencing and T-cell receptor (TCR) data alongside 1x coverage whole genome sequencing from all samples. Two sampling sites were located in Bali province, on the island of Bali, which is directly west of the Wallace Line in central Indonesia, and two in Papua province on New Guinea Island, at the easternmost part of the country and of the Indonesian archipelago (Figure 1A). Balinese samples were collected in Denpasar (DPS, n = 60), the capital and largest city of Bali province and in Pedawa (PDW, n = 52), a village in the mountains of northern Bali known as one of the Bali Aga communities, where the residents preserve ancestral traditional customs and social structures, including long-standing subsistence-level agricultural practices. Papuan samples were collected in two small townships, Sereh (SER, n = 42), located at the base of Mount Cyloop and Yoboi (YOB, n = 45) on the shores of Lake Sentani, both located near Jayapura, the capital of Papua province. The four sites are characterised by distinct environments and lifestyles (Figure 1B). Donors from Denpasar, Pedawa and Yoboi reported that their families had lived in the same place for at least three generations; donors from Sereh were self-reported first and early generation migrant families from the highland interior of New Guinea.

**Figure 1.**
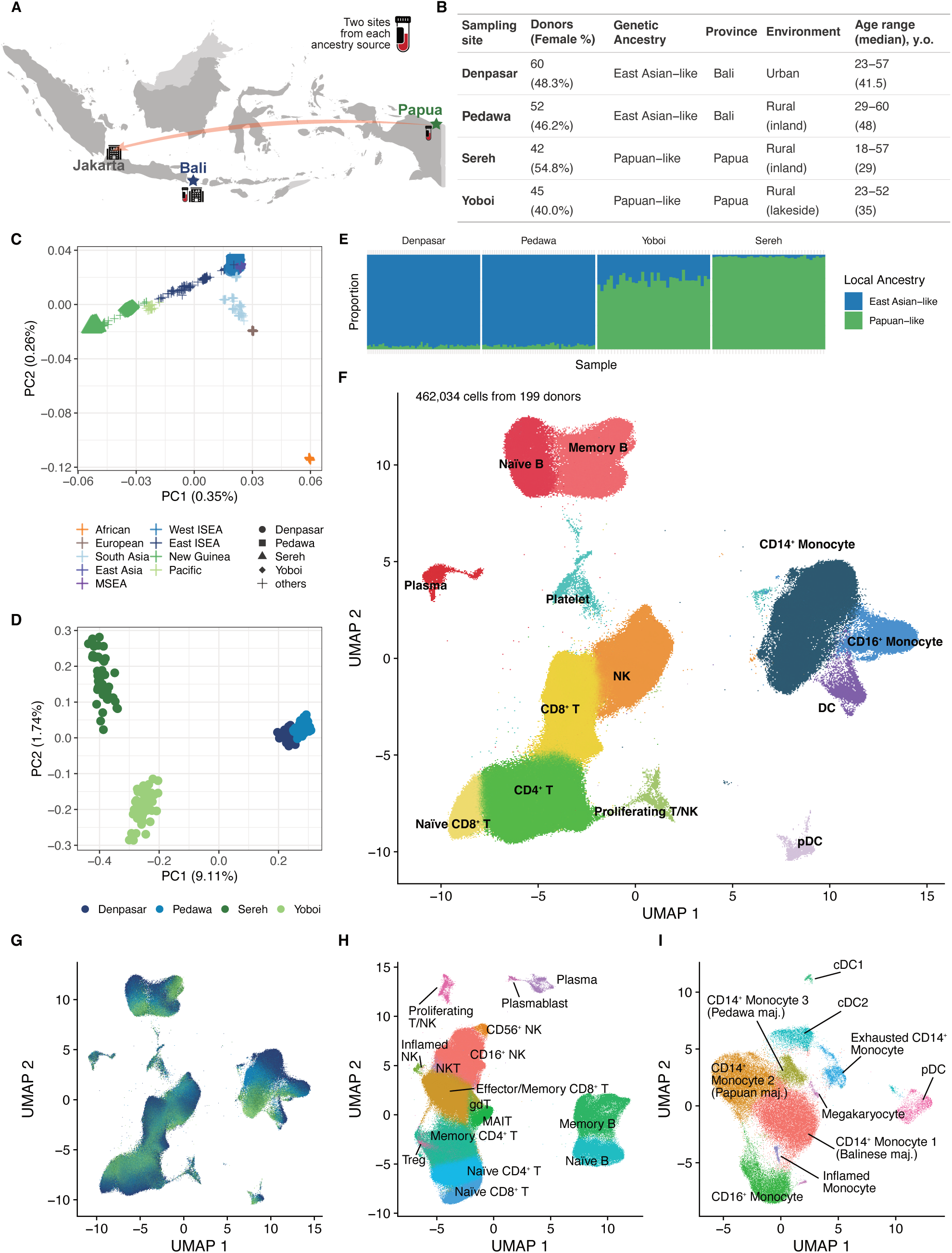
Capturing immune diversity in Indonesia. **A**) Map of Indonesia showing sampling and PBMC isolation sites. Sampling sites in Bali represent urban and inland rural environments, while sites in Papua represent inland and lakeside rural areas. PBMCs from Balinese donors were isolated in Denpasar, Bali, whereas samples from Papuan donors were flown to Jakarta and processed at the Mochtar Riady Institute for Nanotechnology (MRIN). **B)** Donor demographics across the four sampling sites. **C)** Principal component analysis of genotype data for the 199 study sampels alongside samples from the 1000 Genomes Project and external Indonesian & PNG datasets, colored by ancestry. **D)** PCA of genotype data for the 199 study samples, colored by sampling site. **E)** Local ancestry inference of study samples as estimated by RFMix v2. **F-G)** UMAP of PBMCs colored by level 1 cell type annotation (F) and by sampling site (G). **H-I)** UMAP of lymphoid cells (H) and myeloid cells (I), colored by level 2 annotations.

We performed principal component analysis (PCA) of genotype data at the global level to understand the broad genetic affinities of all donors (Figure 1C). Balinese donors from both groups (Denpasar and Pedawa) clustered with other samples from East Asia, Mainland Southeast Asia and West Island Southeast Asia (individuals from Sumatra and its sub-islands, Borneo, Java, and Bali) when examining the first two principal components. The two Papuan groups showed divergent positions, with individuals from Sereh clustering with highland populations from Papua New Guinea, while those from Yoboi were shifted toward groups from eastern Indonesia and clustered together with other comparative samples from the New Guinean Bird’s Head. Principal component analysis of study participants likewise revealed that the first two principal components separate Papuan and Balinese donors, as well as Sereh and Yoboi, although Denpasar and Pedawa do not form independent clusters (Figure 1D). We therefore employ two types of population descriptors to refer to donors, distinguishing between geographic descriptors that indicate where sampling took place and thus capture environmental diversity such as Balinese and Papuan, and genetic similarity descriptors that capture global and regional genetic population structure, such as East Asian-like and Papuan-like.

The F4-ratio test [21] and RFmix local ancestry [22] analyses both indicate that donors from Yoboi harbor around 25% East Asian-like genetic ancestry (∼75% Papuan-like genetic ancestry; Figure 1E), which is also visible in the global admixture proportions (Figure S1A). Additionally, admixture tests detect weak but consistent Papuan-related gene flow into the Balinese, at ∼3% in Denpasar and ∼5% in Pedawa (Figure S1A,B) around ∼1,000 years ago (Figure S1C). MSMC-IM [23] cross-coalescence analysis indicates a deep divergence between Balinese and Papuan populations (*>*20,000 years ago), and a more recent split between Sereh and Yoboi (∼ 9,500 years ago; ∼16,000-5,000 years ago) (Figure S1D). Cross-coalescence for the two Balinese groups did not reach zero, indicating a close affinity or continuing recent gene flow between Denpasar and Pedawa, rather than a complete early separation. Altogether, these analyses confirm that our samples span the characteristic regional East Asian-Papuan-like genetic ancestry cline [2, 24] that characterises the Indonesian archipelago.

scRNA-seq data from all donors was generated across 27 randomised library batches using the 10X genomics Chromium X platform. After initial QC and genetic demultiplexing we integrated the resulting datasets using Harmony. Each library contained at least 20,000 cells (mean = 30,075 cells per library; mean reads per cell = 23,874; mean genes per cell = 1,578; Table S1). The number of cells per donor ranged from 667 to 5,508 (mean = 2,423; median = 2,410; Table S2), for a total of 462,034 cells that passed filtering. After performing graph-based clustering with Seurat and visualizing the cells with UMAP, we manually annotated 13 high-level cell types using canonical gene markers curated from the literature (Figure 1F, Table S3). A majority of cells were annotated as lymphoid cells (385,675), with CD4^+^ T cells (135,461), CD8^+^ T cells (91,629), and NK cells (81,409) the most abundant. We also identified a total of 68,916 myeloid cells, with CD14^+^ monocytes (51,039) being the most abundant. Although each identified cell type formed a distinct cluster, separation driven by the four sampling sites was readily apparent within each cell type (Figure 1G). In-depth manual annotation of the lymphoid and myeloid lineages is shown in Figure 1H–I. Using curated marker gene sets we identified 16 lymphoid subtypes and 10 myeloid subtypes, including two inflamed clusters (NK and monocyte), both of which were heavily dominated by cells from one donor (DPS-006, who was removed from all analyses except eQTL mapping). We also identified distinct separation of CD14^+^ monocytes into three clusters: two of majority-Papuan and majority-Balinese cells, and a smaller cluster with a subset of cells from Pedawa donors further discussed below.

### Disentangling biological and technical variation

PBMC isolation for all Balinese samples took place at Universitas Udayana, in Denpasar (Figure 1A). However, due to limited access to local equipment and facilities, PBMC isolation from Papuan samples took place at the Mochtar Riady Institute for Nanotechnology (MRIN), near Jakarta (3,804 km away by air from Jayapura). This resulted in a technical artefact whereby samples from Papuan donors generally have higher times to PBMC isolation (median = 19 hours) than Balinese donors (median = 5.7 hours). Samples collected in Denpasar were processed the quickest (median = 3.75 hours) and in Sereh the slowest (median = 28.9 hours; Figure 2A). Although all cell type clusters contain cells from donors from all four sites (Figure 2B, Figure 1G), there is significant structuring of clusters by geography. We also observed associations between sampling time and cell type proportions across the dataset, significantly affecting nine of the level 1 cell types (Figure 2C). Taken together, these observations suggested a potential conflation of inter-group biological differences of interest with unwanted technical variation introduced by differences in sample processing.

**Figure 2.**
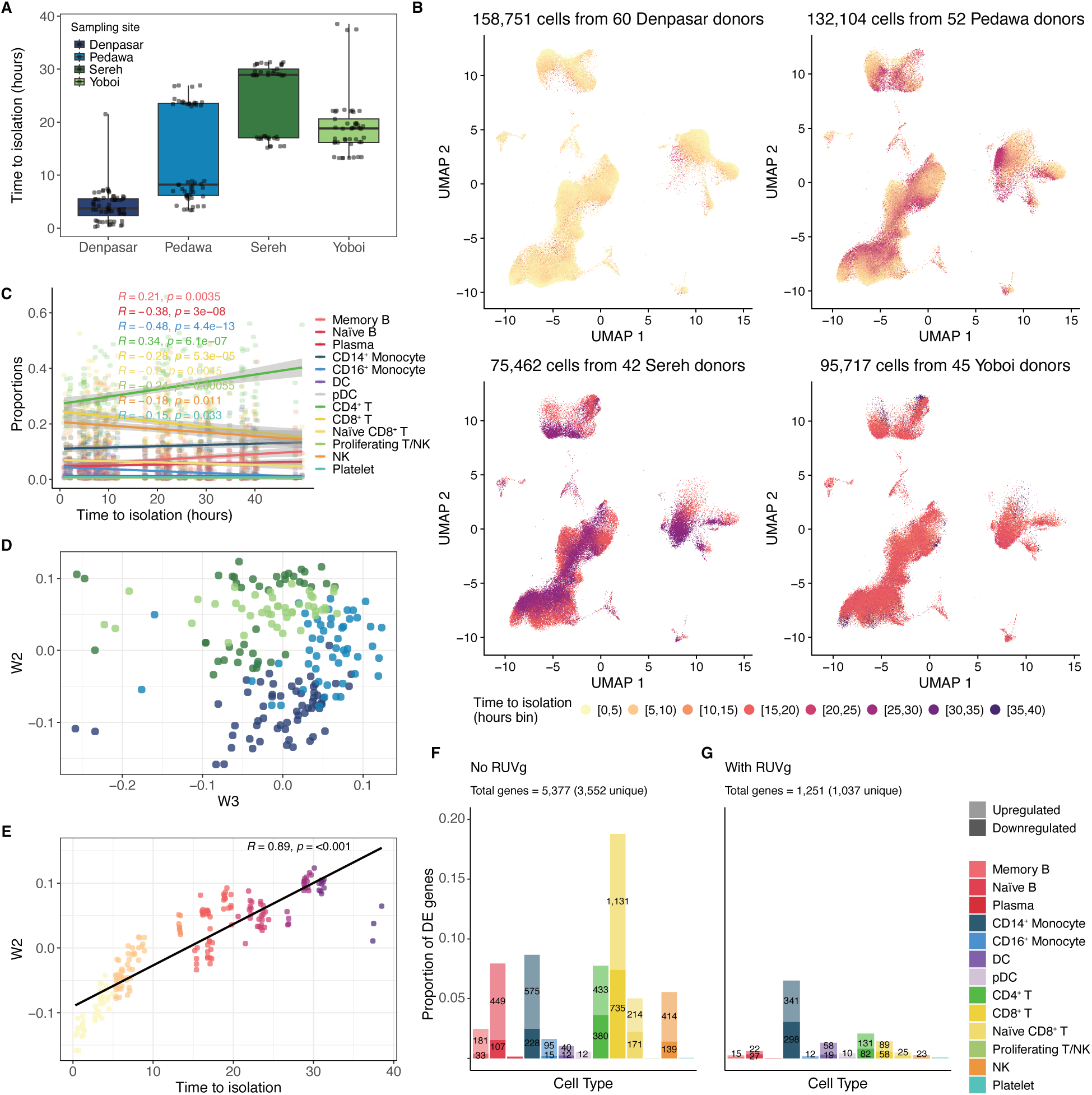
Impact of time to PBMC isolation on cell type proportion and gene expression. **A)** Boxplots showing time to isolation across sampling sites, with each dot representing a sample. **B)** UMAPs of all PBMCs per sampling site, colored by binned time to isolation. **C)** Correlations between time to isolation and unadjusted cell type proportions. **D)** Scatter plots of factors of unwanted variations (*Ws*) 2 and 3 (*W* 2 and *W* 3) from RUVg colored by sampling site. **E)** Pearson’s correlation between *W* 2 and time to isolation. **F-G)** Bar plots of differentially expressed genes (F) before and (G) after adjusting for *Ws* per cell type. Age, sex, sequencing batch, time to isolation, and cell counts were also included as covariates.

We explored this possibility using RUVg [25], a modeling framework that removes unwanted variation from pseudobulked expression data based on 1,124 stably-expressed genes validated on PBMC data [26]. RUVg was able to detect multiple linear factors in our data (termed *Ws*) that captured variation between sampling sites (Figure 2D). *W2*, in particular, correlated significantly with time to isolation (*p* = 1.84 × 10*^−^*^69^; Figure 2E). To determine a suitable W value for our dataset we took advantage of the fact that the times to isolation for samples from Pedawa and Sereh were bimodally distributed (Figure 2A). As all donors reported being healthy at the time of collection, differential expression between samples from the same site that were processed early or late is primarily attributable to technical confounders.

DE testing between Pedawa early and late samples using a model that included age, sex, sequencing batch, cell count, and time to isolation but no RUVg *Ws* as covariates identified 3,552 unique DE genes in at least one cell type. This effect was most pronounced in CD8^+^ T cells (n = 1,866, 18.78% of all tested genes), CD14^+^ monocytes (n = 803, 8.66%) and naïve B cells (n = 556, 7.93%; Figure 2F). We then repeated DE testing including increasing numbers of *Ws* identified by RUVg as additional covariates. At *W* = 8, overall unique DE genes between Pedawa early and late fell to 1,037, with a majority of signal coming from CD14^+^ monocytes (n = 639, 6.5% of all tested genes); Figure 2G) and very low numbers of DE genes in other cell types. We observed similar trends when comparing between early and late Sereh samples (Figure S2D–E). Since the Pedawa-specific CD14^+^ monocyte cluster we identified in Figure 1I contained primarily samples from Pedawa late donors, we chose to include 8 *Ws* as covariates in all expression analyses downstream, as it struck the balance between removing unwanted variation and retaining biological signal.

### Expression quantitative trait loci (eQTL) analysis reveals cell type-specific differences tied to genetic structure

We performed eQTL analysis using a linear mixed model implemented in quasar [28], which allowed us to control for local kinship structure present in our data (Table S4). In this analysis we considered only genes expressed in at least 10% of donors in nine different cell types, excluding pDCs, plasma cells, platelets and proliferating T/NK cells due to generally low cell counts across individuals. Instead of relying on two-pass multiple testing correction, we used ACAT [29] to calculate gene-level p-values; this allowed us to identify significant eQTLs in 2,835 genes across the dataset at a false discovery rate (FDR) of 5%, ranging from 1,539 in CD4^+^ T cells to 155 in DCs (Table S5). Only 24 genes were significant across all cell types, with any given eQTL being detected in an average of 2.12 cell types; 1,423 (50.19%) of genes had a significant eQTL in only one cell type. We then applied Bayesian shrinkage to the quasar output using *mashr* [30], leveraging information across cell types to increase our discovery power. At a threshold of *LF SR <* 0.05 we identified significant eQTLs in 5,384 genes across all nine cell types (Table S5).

As expected, mashr led to an overall increase in the number of eGenes identified in each cell type, with an average of 4,181 eQTLs per cell type and a near-majority of eQTLs (n = 2,686; 49.89%) being shared across all nine cell types when considering statistical significance only (*LF SR <* 0.05 in both cell types; Figure 3A). This higher eQTL sharing between more immunologically similar cell types can also be seen when we consider a statistically significant eQTL shared with a second cell type if the estimated effect size (*β*) in the two cell types is within a factor of 0.5 (Figure 3B).

**Figure 3.**
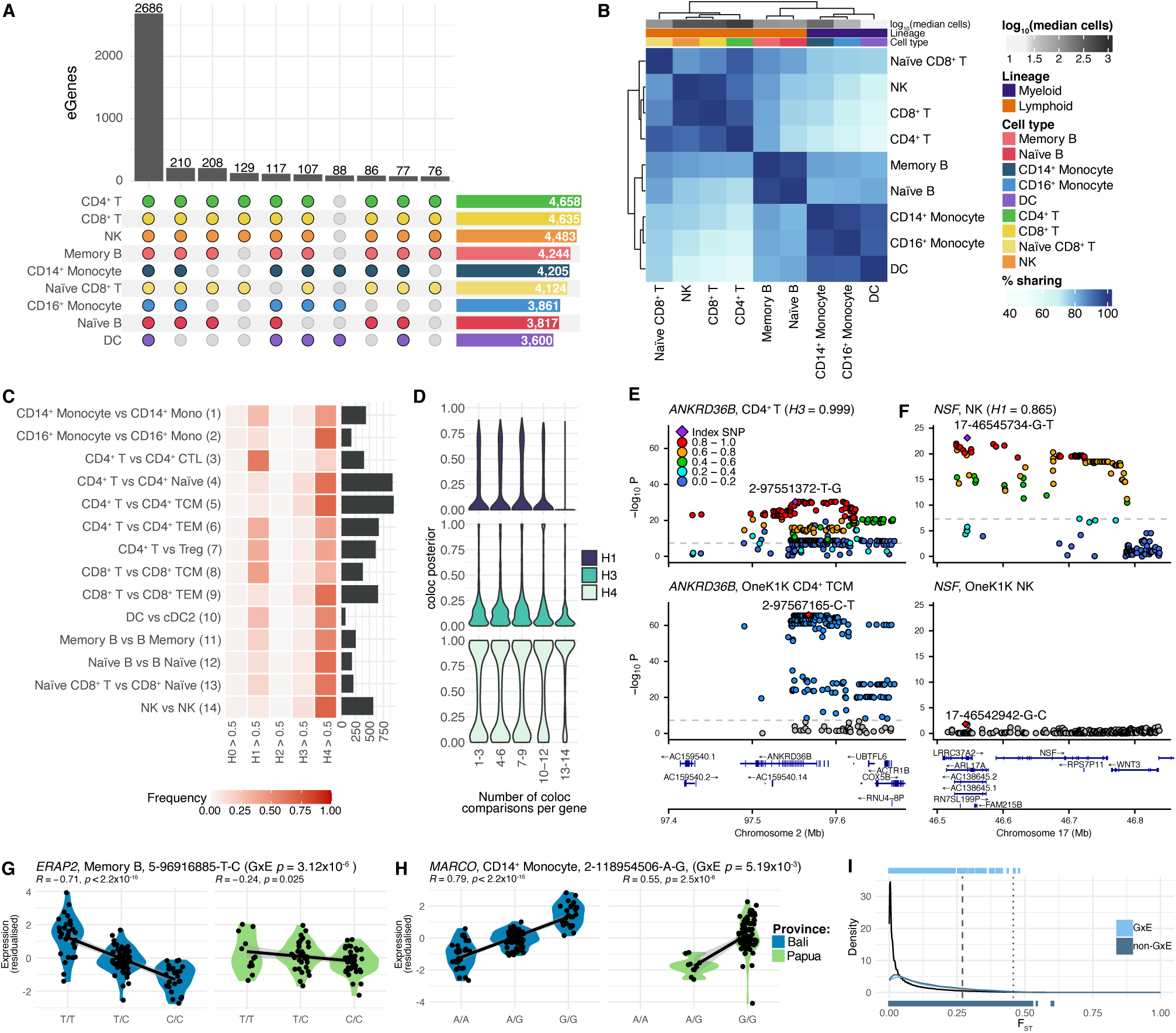
eQTL mapping and genetic effects on gene expression across Indonesia. **A)** Upset plot showing number of significant (*LFSR <* 0.05) eGenes after mashr across cell types. Only the 10 largest categories are shown. **B)** Significant eGene sharing between tested cell types based on effect size (*β*) ratio after mashr. **C)** Colocalisation of eQTLs from this study with their closest OneK1K cell type. OneK1K cell type labels as in the eQTL Catalogue. The number of genes considered for each comparison is indicated by the bar plot. Definitions of *H*0 to *H*4 are as in [27]. **D)** Distributions of *H*1, *H*3 and *H*4 posterior probabilities across all eGenes tested for colocalisation between Indonesia and OneK1K, stratified by the number of pairwise comparisons an eGene was involved in. **E**-**F)** Examples of colocalisation against OneK1K data. Indonesian data is shown on the top and OneK1K on the bottom for **E)** *ANKRD36B* in CD4^+^ T cells and **F)** *NSF* in NK cells. Pairwise LD is not available for OneK1K; instead, red indicates the index SNP and blue points exceed nominal genome-wide significance. **G**-**H)** Residualised expression across the two Indonesian provinces in the data for genes with a significant GxE interaction term. Expression is shown relative to genotype at the most significant eSNP across all of Indonesia for **G)** *ERAP2* in memory B cells and **H)** *MARCO* in CD14^+^ monocytes. **I)** Distributions of F_st_ values genome-wide and across all lead eSNPs in the dataset. The dashed line indicates the empirical top 5% of the genome-wide distribution, the dotted line the empirical top 1%. Rug plots indicate F_st_ values for individual eSNPs, separated by the presence or absence of a significant GxE interaction.

To investigate how global genetic population structure influences eQTL detection, we performed colocalisation analyses [27] between our Indonesian data and eQTLs mapped in OneK1K, a cohort of 982 individuals of Northern European ancestry [31] that represents one of the largest publicly available scRNA eQTL datasets from whole blood. Since the OneK1K data was not processed using mashr we used the quasar test statistics and considered only significant Indonesian eQTLs in this analysis, further filtering to those with at least one eSNP with *p <* 10^4^ (n = 1,728, see methods). We performed colocalisation on each comparable cell type from OneK1K’s more granular annotation, resulting in a total of 14 comparisons (Table S6). We define an eQTL as colocalised between populations when *H*4, the posterior probability of them sharing a single causal variant is ≥ 0.5. On average, most eQTLs shared a causal variant in the two cohorts (mean % of colocalised genes = 51.36%; Figure 3C). The different levels of annotation and resolution between the two cohorts likely influence this result, as the lowest colocalisation fraction is found when comparing Indonesian CD4^+^ T cells and OneK1K’s CD4^+^ T CTL (16.17%), a rare CD4^+^ subtype. Any cell type-specific eQTLs detected by OneK1K at more granular resolutions than those captured by our annotation are likely to be dampened or absent in our dataset.

We also examined eGenes where *H*1 or *H*3 *>* 0.5, which respectively represent the posterior probability that there is an eQTL only in Indonesia, or that there are eQTLs in both populations but with different causal variants (Figure 3C). Across all cell types *H*1 exceeded 0.5 an average of 22.83% of the time, although again this number varied greatly, from 60.30% of the time when comparing between Indonesian CD4^+^ T cells and OneK1K’s CD4^+^ T CTL, to 6.65% when comparing Indonesian CD4^+^ T cells to OneK1K’s CD4^+^ T TCM, which are direct equivalents of each other (Table S6). *H*3 was > 0.5 between 1.71% (CD16^+^ monocytes) and 12.43% of the time (Indonesian CD4^+^ T cells and CD4^+^ TCM), with an average of 7.13% of genes per pairwise comparison. Nevertheless, eQTLs that failed to colocalise in one cell type often colocalised in other, closely related ones. Indeed, we observed a general association between the number of cell type comparisons we performed coloc in and the relative likelihood of colocalisation (*H*4) versus non-colocalisation (*H*1 or *H*3 *>* 0.5; Figure 3D, Figure S3A). Specifically, as the number of times we performed colocaliation for a specific eGene increased, the log odds ratio significantly favored the shared causality (*H*4) hypothesis (p = 9.13 × 10*^−^*^6^). We also observed a significant association between individual posterior probabilities and the number of coloc comparisons an eGene was included in: individual *H*4 values significantly increased (*p <* 2 × 10*^−^*^16^) as the number of comparisons a gene was involved in increased, while *H*1, but not *H*3, decreased (*H*1*p* = 4.2 × 10*^−^*^9^; *H*3*p* = 0.392). These results suggest that genuine populations-specific eQTLs are more likely to impact one or a limited number of cell types. Examples include a tissue-specific eQTL affecting *UCP2*, a widely expressed mitochondrial protein thought to help dampen mitochondrial ROS levels [32], which colocalises between Indonesia and OneK1K in 11 out of 13 comparisons (*H*4 *>* 0.95), but not in NK cells (*H*3 = 0.688; Figure S3B-C), where fine-mapped 95% credible sets contain no SNPs in common.

Additionally, a non-negligible number of genes (141, 11.5% of genes tested at least twice) rarely or never colocalised (Figure S3D-E). Examining these genes can reveal broader instances of population-specific gene regulation. For example, across 10 cell type pairs *ANKRD36B* was consistently associated with *H*3 values *>* 0.5 (*>* 0.99 in 9 comparisons; Figure 3G, Figure S3D). Fine mapping in CD4^+^ T cells identified a 95% credible set containing 27 SNPs, 13 of which were in perfect linkage disequilibrium with each other and have minor allele frequencies (MAF) < 0.02 in all gnomAD population supergroupings except for East Asia, where the average MAF is 0.11. Similarly, *H*1 for *NSF* ranged between 0.515 and 0.865 across all 14 pairwise comparisons (Figure 3H, Figure S3D). In this case, the fine mapped 95% credible set in CD4^+^ T cells contained 16 SNPs, all with appreciable MAF in gnomAD, but there is no significant eQTL reported in any of the OneK1K cell types.

Finally, we tested for gene by environment (GxE) interactions between Bali and Papua to identify instances of differential regulation between the two islands. Across the entire set of eGenes identified by quasar we identified 339 eGenes where this interaction term was significant (Table S7). Results per cell type ranged from 109 significant eGenes in CD4^+^ T or CD14 Monocytes (7.08% and 12.75% of all eGenes in those cell types) to 7 genes (4.52%) in DC, consistent with the lower number of eQTLs discovered in less frequent cell types.

We also identified genes that exhibited subtler trends. For instance, GxE interactions affect expression of *ERAP2* in memory B cells (FDR-adjusted interaction *p* = 3.12 × 10*^−^*^6^) and four other cell types (FDR-adjusted interaction *p* between 0.013 and 0.0012). Although *ERAP2* eQTLs were consistently colocalised between Indonesia and OneK1K (Figure 3E), their effects were greatly attenuated in Papuan donors compared to Balinese (Figure 3G). *ERAP2* is involved in peptide trimming in the ER lumen as part of MHC class I antigen presentation and more broadly in the activation of the immune response, and has been repeatedly argued to be targeted by balancing and positive selection in human populations [33, 34, 35], and of possible adaptive introgression from archaic hominins [36]. In the best studied example, rs2248374, which has a global allele frequency of 0.519 in gnomAD, disrupts a canonical splice site and gives rise to an alternative transcript that is degraded by nonsense-mediated decay, ultimately leading to a decrease in MHC Class I receptors being expressed on the surface of B cells [33]. Although this variant is contained within the 95% credible set in memory B cells in our data, it has a relatively small posterior probability of being causal in our data (0.0005) and is not in the 95% credible set in OneK1K, suggesting the eQTL signal we observe is independent of aberrant splicing, as is its dampening in Papuans.

A more striking example is the eQTL affecting *MARCO*, a pattern recognition receptor expressed by CD14^+^ and CD16^+^ monocytes that mediates the recognition of multiple bacterial pathogens, including *Mycobacterium tuberculosis* [37]. MARCO was strongly differentially expressed between western and eastern Indonesian populations in whole blood bulk RNA-seq data [5]. Here we report an eQTL that colocalises strongly with OneK1K (CD14^+^ monocytes *H*4 = 0.996). Evidence for GxE between Bali and Papua in CD14^+^ monocytes is strong (FDR-adjusted interaction *p* = 0.005), with different expression levels in Bali and Papua (Figure 3H). Genetic variation at the locus is highly structured, with no observed homozygotes for the reference allele in Papuans and high differentiation between the two provinces (Weir & Cockerham’s F_st_ = 0.401, empirical *p* = 0.017).

On the basis of this observation we calculated F_st_ between Papua and Bali for all lead eSNPs in the set of quasar eQTLs (4,931 eSNPs). We observe a general excess of lead eSNPs with F_st_ values in the top empirical 5% relative to the genome-wide distribution (n = 552; binomial *p* = 1.3 × 10*^−^*^67^, Figure 3I). However, a high level of population differentiation is not sufficient to give rise to significant GxE interactions, with eSNPs with and without support for GxE interactions being equally likely to be in the top 5% of the empirical F_st_ distribution (*χ*^2^ *p* = 1). For instance, we observe a similar trend for the phospholipase *PLAAT3* (F_st_ = 0.415, empirical *p* = 0.015; FDR-adjusted interaction *p* = 0.004), which is required for picornavirus infection of human hosts and enables the transportation of the viral genome into the host cytoplasm [38] but not for *SIGLEC10* (F_st_ = 0.520, empirical *p* = 0.005; FDR-adjusted interaction *p* = 0.416), an inhibitory cell-surface receptor that senses foreign glycans [39]. Taken together, these observations suggest that genetic differences between the two sites are only partially responsible for these trends, and pointing towards a substantial contribution from environmental variation in driving this effect.

### Contributions of local ancestry and introgression to gene expression

To investigate how ancestry background shapes regulatory variation in Indonesian populations, we tested whether significant eSNP alleles were correlated with the local ancestry state (LA; determined by RFMix), considering all significant lead eSNPs following mashr analysis (n = 14,896 SNPs). Loci with strong correlations between genotype and LA and/or archaic ancestry indicate that regulatory variation is at least in part carried on ancestry-specific haplotypes and that population history (such as population admixture) has contributed to these differences [2, 3]. Using an *r*^2^ threshold of 0.5 to define variants associated with LA, we identified 139 eQTLs across 9 cell types (see Methods), ranging from 13 eQTLs in Naive B cells to 18 eQTLs in CD8^+^ T cells (Figure 4A). The majority of these LA-associated alleles are more frequent in Papuan rather than Balinese population (Figure 4A). This is consistent with ancestry-differentiated allele frequencies driving a portion of regulatory differences between groups. Several of top LA-associated eGenes (*r*^2^ > 0.7, showing consistent correlation between genotype dosage and ancestral dosage (Figure 4B-C), Table S8) have immune roles, such as *SLC35A1*, a Golgi sialic-acid transporter that is essential for Influenza-A virus receptor expression and viral entry [40]; *CST7* which encodes Cystatin-F, a cystein-protease inhibitor which is highly expressed in cytotoxic lymphocytes where it modulates granzyme activation and NK cell effector function [41]; or *CDIP1*, which plays a role in TNF-*α*-mediated apoptotic signaling which mediates the inflammatory cell-death pathways [42]. These results show that ancestry-specific haplotypes arising from recent admixture may contribute in shaping the gene regulation in the populations.

**Figure 4.**
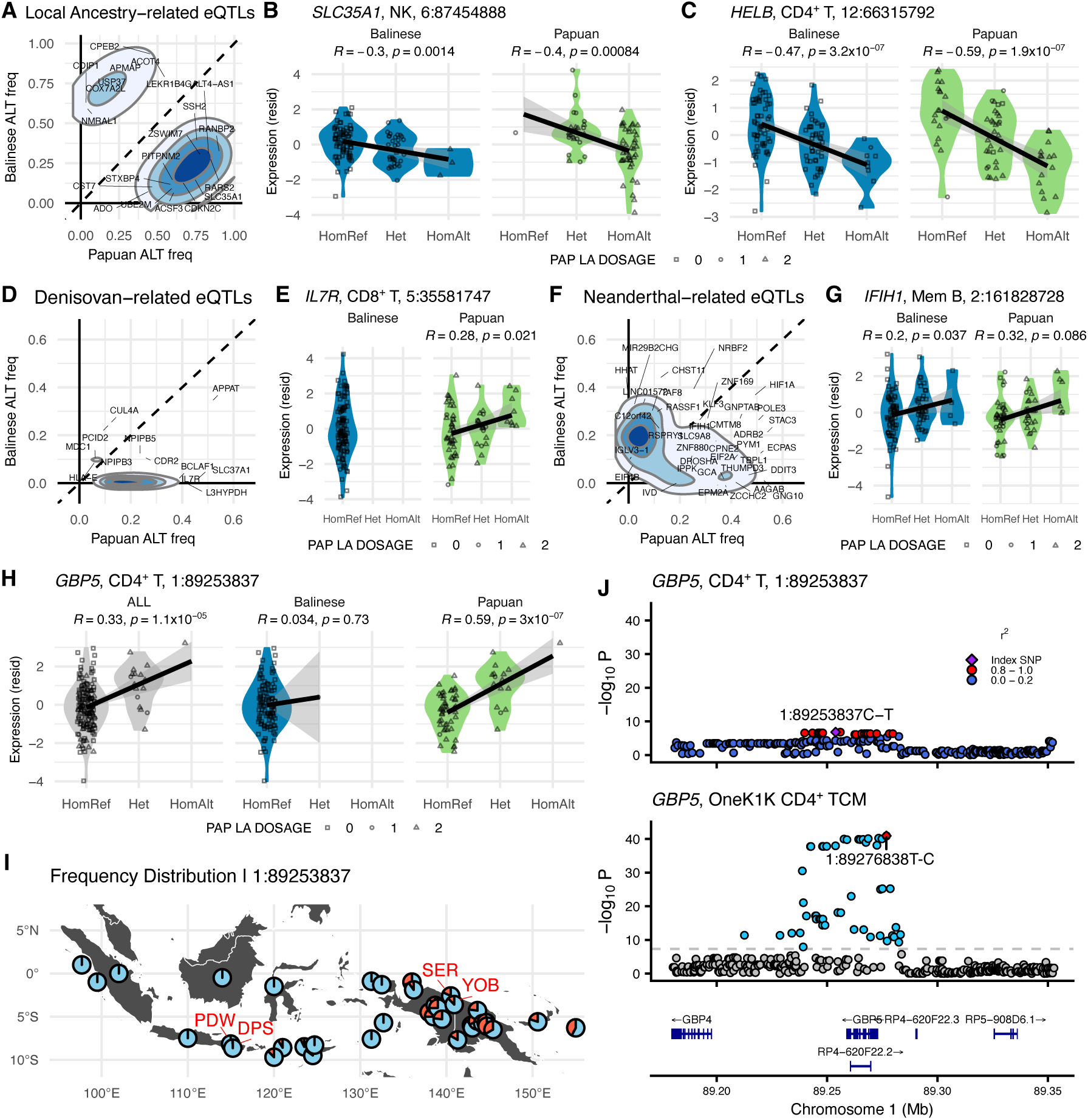
Contribution of local ancestry and archaic introgression to regulatory variation. **A**) Distribution of allele frequencies for LA-associated eQTLs in Balinese and Papuan samples. **B)** Example of a strong LA-associated eQTL (*SLC35A1*): a violin plot of genotype dosage vs. residual expression; lead SNP has *r*^2^ = 0.79. **C)** Example of a weak LA-associated eQTL (*HELB*): same plot format as B, lead SNP *r*^2^ = 0.02. **D)** Distribution of allele frequencies for Denisovan-associated eQTLs in Balinese and Papuan samples, showing alleles at moderate frequency in Papuans and near zero frequency in Bali. **E)** Violin plot of an example Denisovan-associated *IL7R* eGene. **F)** Distribution of allele frequencies for Neanderthal-associated eQTLs in Balinese and Papuan samples. **G)** Violin plot of an example Neanderthal-associated *IFIH1* eGene. **H)** Violin plot showing genotype-expression on the putative Denisovan-introgressed eQTL at 1:89253837 for *GBP5*. The eSNP shows the HomAlt class absent in Balinese. **I)** Geographic scatterpie plot of allele frequency at 1:89253837 across Indonesian and Papua New Guinean populations [3, 6, 10]. Each pie indicates ALT allele frequency in red and REF allele frequency in blue. **J)** LocusZoom-style plots centered at 1:89253837 for *GBP5* in this dataset (top) and OneK1K’s index SNP (bottom), with the LD *r*^2^ value categories shown for this dataset.

At deeper evolutionary timescale, introgression from archaic hominins has also introduced population-specific variations into the genomes of modern populations, and asked whether Denisovan– and Neanderthal-derived haplotypes also harbor variants that influence gene expression. We began by identifying individual-level introgressed haplotypes using HMMix [43], validating the accuracy of these calls against high-coverage genomes that also had available 30x whole-genome sequencing data. The inferred fragment length overlapped by an average of 83% (Figure S4A-B), which shows concordance of our low– and high-coverage inferences. In our dataset, archaic introgression covered an average of 1.7% of the genome per individual, peaking at 3.1% and 3.6% in Yoboi and Sereh, respectively. We observed the most Denisovan-introgressed segments in Sereh (91 Mb per genome), then in the Yoboi (68 Mb), followed by the Balinese groups (5 Mb) (Figure S4C-D). In contrast, the inferred Neanderthal introgressed segments were fairly consistent across our sample groups (73–77 Mb) (Figure S4D). The pattern of high Denisovan coverage in Papuan highland and lowland groups and low Denisovan coverage in Bali is consistent with prior work showing that elevated Denisovan ancestry is common in Papuan populations and rare elsewhere [3].

Focusing on variants on Denisovan-introgressed haplotypes, we found 179 eQTLs across 9 cell types, ranging from 15 eQTLs in both CD4^+^ T and Naive CD8^+^ T cells to 24 eQTLs in CD14^+^ Monocytes. A majority of the Denisovan alleles that underlie these signals occur at moderate-to-high frequency in Papuans and are very rare in Bali (Figure 4D). eGenes with robust eSNP introgression signals include *IL7R*, *STAT2*, or *HLA-E* (93-100% introgressed-haplotype ALT frequency) (Figure 4E, Figure S4E, Table S9). The lead eSNPs associated with these signals have some of the highest introgressed allele frequencies in the Papuan group in dataset (*X* = 21%; median = 20%), compared to the Balinese group (*X* = 5%; median = 0.9%). These genes contribute to T-cell homeostasis [44], type I interferon antiviral signalling [45], and NK-cell regulation [46], respectively. At lower allele frequencies we find a strong signal of Denisovan introgression driving an eQTL affecting *SMAD1* levels, a gene that regulates BMP/SMAD-dependent control of inflammatory signaling [47]. This suggests that Denisovan introgression may impact immune regulation in these populations. Furthermore, we identified 370 eQTLs across 9 cell types that are associated with Neanderthal introgression, ranging from 33 eQTLs in Naive B to 36 eQTLs in Memory B (Figure 4F). As above, several introgression-associated genes are involved in immune defense. *IFIH1* (97% introgressed haplotypes carrying ALT) (Figure 4G; Figure S4F) encodes the cytosolic RNA sensor MDA5, a receptor that detects viral double-stranded RNA and stimulates type-I interferon response [48], while *TLR1* is a known target of Neanderthal introgression and part of the Toll-like receptor complex that recognizes bacterial triacylated lipopeptides at the cell surface [49].

All available Denisovan genome sequences are derived from samples found in Denisova cave, roughly 8,000 km away from Jayapura. Their high divergence from the introgressing Denisovan population can make confident inference of introgressed Denisovan ancestry challenging, especially in Papua [50, 3, 16]. An example of this is an eQTL impacting *GBP5*, an interferon-inducible GTPase that promotes NLRP3 inflammasome activation and restricts enveloped viruses [51]. Approximately 13% of Papuan individuals in our dataset carry overlapped introgressed haplotypes that spans *GBP1* to *GBP6*, these haplotypes have been previously thought to be a target of adaptive introgression in an independent cohort of individuals of Papuan-like genetic ancestry [50]. However, the sequence of some of these haplotypes, particularly the *GBP5*, is as close to that of both Vindija and Chagyrskaya Neanderthal as to the Altai Denisova reference, such that we cannot confidently assign a specific source to it. Yet at the lead eSNP, 1-89253837-C-T, for this gene, all but one Balinese individual are fixed for the ancestral allele (Figure 4H), and there are only 19 observations of the alternative allele worldwide reported in gnomAD. Amongst Papuan individuals in our dataset this same allele has a frequency of [ 18%] and is clearly associated with differences in expression levels (Figure 4H); other Papuan, Eastern Indonesian, and Oceanian populations [3, 6, 10]—all of whom are also absent from gnomAD—also carry this allele at appreciable frequencies (Figure 4I). Examination of LD structure around the region reveals that the signal is driven by a set of 18 SNPs between which pairwise LD never drops below 0.8, but that are not in high LD with other variants segregating in the region, a signal that is absent from OneK1K (Figure 4J). The combination of extremely low global allele frequencies with this extended linkage pattern aligns with the properties and distribution of confidently identified Denisovan-related alleles, which only occur in moderate-to-high frequency in Papuan and Papuan-admixed populations [3, 16], and identify the eQTL on *GBP5* and its extended haplotype as likely introgressed from Denisovans. Overall, our results suggest that both Denisovan and Neanderthal introgression contributed functionally relevant alleles to pathways important for immune adaptation to Papuan individuals, and match previous work that showed Denisovan ancestry in Papuans enhances immune functions and gene regulation [52].

### Co-expression networks show island and village –specific patterns in immune regulation

We purposefully designed our sampling strategy to simultaneously capture diversity along genetic ancestry and environmental axes by including participants from two communities with different lifestyles and local environments within each province. To explore this further, we performed co-expression network analysis to investigate groups of genes with similar expression patterns, reflecting shared or distinct functions or biological pathways across our cohort. We used the hdWGCNA R package to construct co-expression networks for five cell types with highest average expression and cell counts, and identified a total of 279 statistically significant (*FDR* ≤ 0.05) modules across all cell type/site combinations (Table S10). To examine whether modules from one group shared similar patterns with those from another group, we conducted module preservation analysis using NetRep. Figure 5A summarizes the number of modules per group and cell type and their preservation status. Across all cell types and sites, 175 modules (62.7%) were seen in only one site, followed by 61 modules (21.9%) that were preserved across all sites. Summed across the five cell types tested we identified the smallest number of modules in Denpasar (48, 26 of which were private to this site, binomial *p* = 0.142), and the greatest in Yoboi (86, 60 of them private, binomial *p* = 0.931). But patterns of module preservation varied across sites. For example, 10 modules identified in Denpasar (21%), such as CD8^+^ T Denpasar-4 and NK Denpasar-4 were replicated in at least one of the two Papuan sites. However, only 6 modules (10%) identified in Pedawa were shared with at least one of the Papuan sites. but not in the other Balinese site, Pedawa. Likewise, across all cell types only 2 modules were private to Bali, but 9 were common to the two Papuan sites. Shared and common modules also tended to have different biological functions. Enrichment analysis indicated that fully preserved modules were mainly enriched for general cellular programs, including cell cycle and proliferation-associated, such as MYC targets v1 (Figure 5B, Table S11), non-preserved modules were associated with diverse functional pathways and immune roles (Table S12). Only a small number of broadly replicated modules, such as NK Pedawa-18 and CD4^+^ T Yoboi-4, were linked to immune response pathways, including interferon signaling.

**Figure 5.**
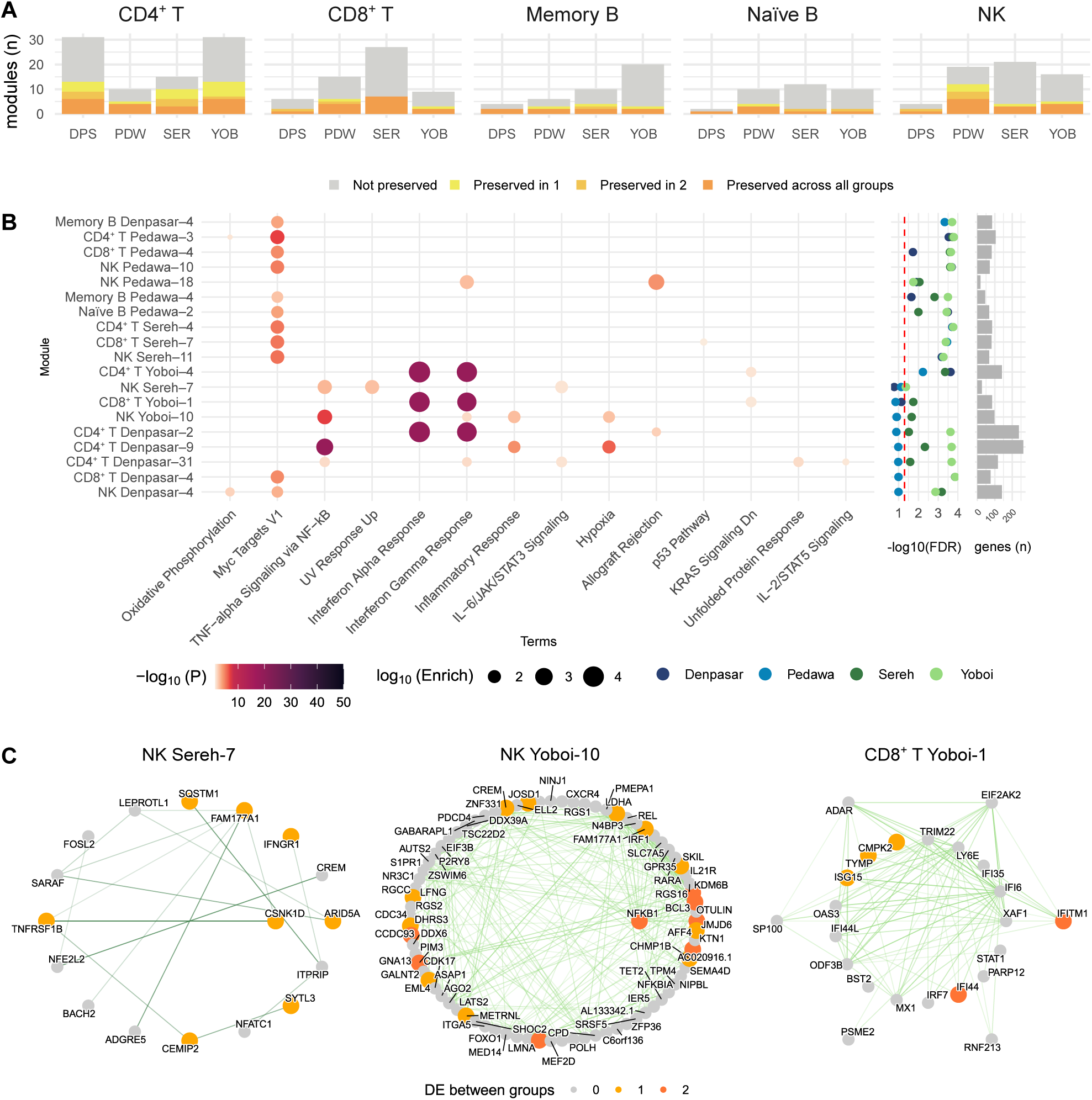
Gene expression analysis showing viral-related pathway associations in specific cell types across two populations. **A**) Bar plot summarizes the number of co-expression network modules identified per site and cell type. Bars are colored by preservation category, indicating whether each module is not preserved, preserved in one or two other sites, or preserved across all sites. **B)** Significant modules identified across five cell types (CD4^+^ T cells, CD8^+^ T cells, NK cells, naïve B cells, and memory B cells), with functional enrichment assessed using MSigDB Hallmark 2020 gene sets. The color of the dot denotes –log(p-value) and size represents the enrichment score. The following panel displays module preservation across sites, with dots colored by site and positioned according to the average adjusted FDR from NetRep. The red dashed line marks –log(FDR) = 0.05. The rightmost bar plot indicates the number of genes contained in each module. **C)** Network plots of Papuan modules that were preserved only within island. Node color indicates genes that are differentially expressed across groups, centrally located nodes represent hub genes.

Finally, we identified hub genes (*kME* ≥ 0.8) for all networks, with three examples of Papuan-only networks shown in Figure 5C. NK Sereh-7 (containing 26 genes)and associated with TNF-alpha signaling via NF-kB, showed *CSNK1D* as the hub gene. NK Yoboi-10 (98 genes) was also enriched for TNF-alpha signaling via NF-kB and inflammatory response, and had *NFKB1* as its sole hub gene. Finally, CD8^+^ T Yoboi-1 (84 genes), which was associated with immune response pathways such as interferon alpha and gamma response, included *CMPK2*, *OAS3*, *ISG15*, and *IFI44* as hub genes, along with *IFITM1*, which is also involved in immune function [53, 54, 55, 56]. Consistent with the results here, differential expression testing (Table S13) revealed that *NFKB1, IFITM1, ISG15, CMPK2*, and *IFI44* were upregulated in Yoboi compared to other groups, and *CSNK1D* was upregulated in the Papuan groups relative to the Balinese groups.

### Clonal expansion in T cells and viral-related transcriptional signatures

To investigate how the local environment shapes adaptive immunity, we analysed T-cell receptor (TCR) diversity in our cohort. After quality control, which included removal of low-read cells and exclusion of orphan clones, we retained 119,527 cells across 198 individuals with complete and high-quality TCR sequences for downstream analysis. Site-specific averages of 921 cells in Denpasar, 539 cells in Pedawa, 404 cells in Sereh, and 349 cells in Yoboi correlated with PBMC isolation time, as reflected in both the number of unique clonotypes and Hill’s diversity (Figure S5A-C). To explore how demographic metadata relate to TCR repertoire diversity across individuals, we examined associations of clonal diversity with age, sex, BMI, and sites across level 2 lymphoid annotations. In naïve CD8^+^ T cells and MAIT cells, clonal diversity decreased with increasing donor age (Figure S5E), replicating well-established trends [57]. Female donors exhibited higher clonal diversity in memory CD4^+^ T cells and MAIT cells (Figure S5F). Additionally, donors with higher body mass index (BMI) tended to show a higher Gini coefficient, indicating greater clonal expansion (Figure S5G). Increased CD4^+^ T cell counts are associated with obesity [58], as adipocytes can directly activate CD4^+^ T cells through HLA class II and leptin, promoting inflammatory response [59, 60].

We then examined clonal expansion dynamics across all samples. Within each individual we labeled clonotypes present at *>* 1.5× the median frequency as expanded. Expanded clones were observed in proliferating T/NK, effector/memory CD8^+^ T, MAIT, memory CD4^+^ T, and NKT cells(Figure 6A-B). We computed the Gini coefficient for each individual and cell type, and tested for differences in coefficient values across sites after correcting for age, sex, and time to isolation. Effector/memory CD8^+^ T cells were the only cell types at level 2 annotation that showed significant differences in Gini coefficient between sites (ANOVA *p* = 0.016; Figure 6C), with posthoc analyses showing that the main driver of this effect were differences between Denpasar and Yoboi donors.

**Figure 6.**
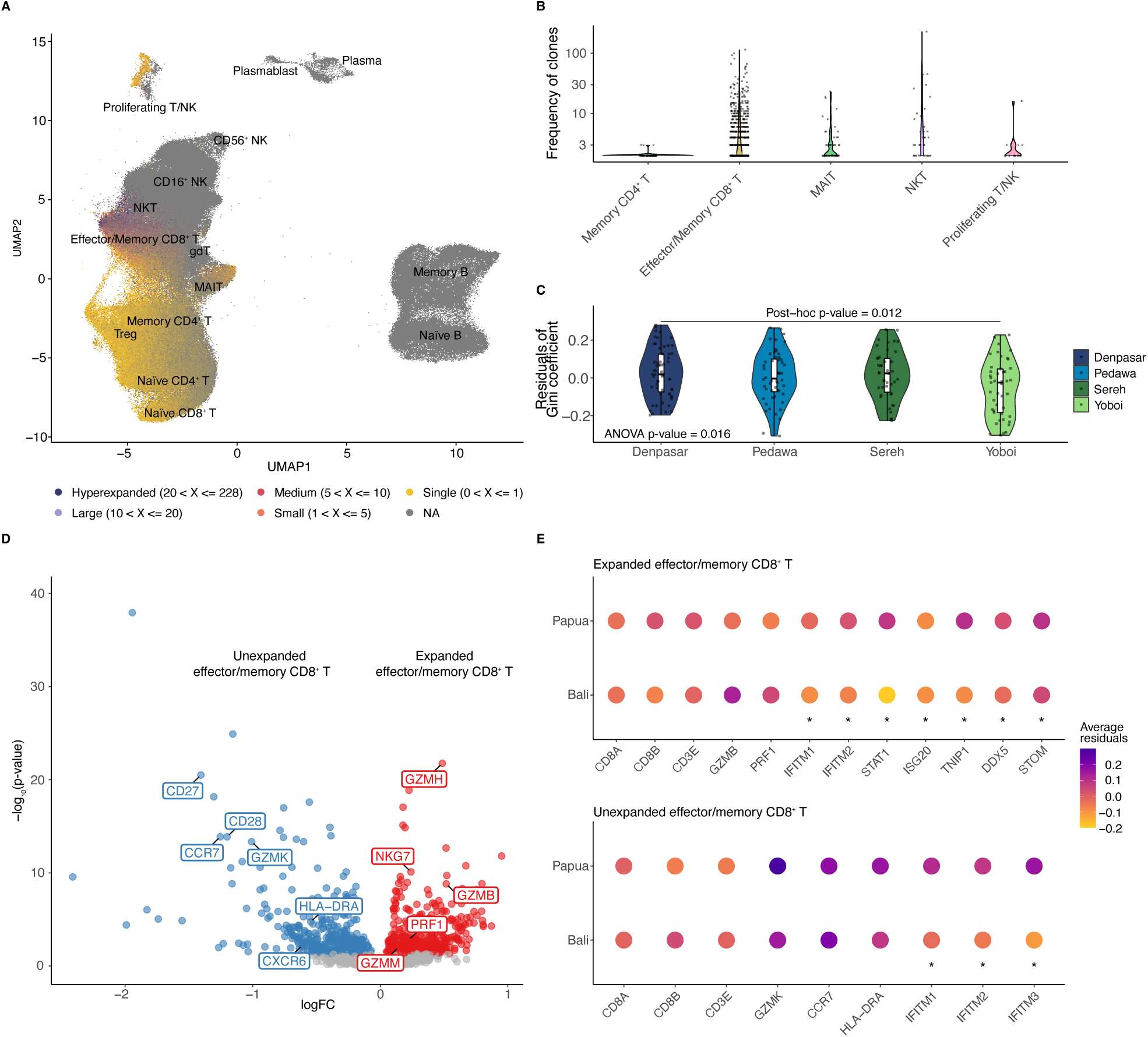
Clonal expansion and interferon-stimulated gene signatures in effector/memory CD8^+^ T cells. **A**) UMAP visualisation of lymphoid cells labeled by clonal size. Colours indicate clone size categories: NA (cells without complete TCR), single (1 clone), small (2-5 clones), medium (6-10 clones), large (11-20 clones), and hyperexpanded (21-228 clones). Cell type annotation level 2 is overlaid on the UMAP. **B)** Violin plot showing the frequency of clones in each level 2 cell type. Only cell types containing expanded cells are shown. **C)** Residuals of Gini coefficient in effector/memory CD8^+^ T cells across the four sampling sites after adjusting by age, sex, and time to isolation. Residuals represent clonal expansion with significance tested by one-way ANOVA. **D)** Volcano plot of differentially expressed genes between expanded and unexpanded effector/memory CD8^+^ T cells. Log fold-change of the average expression were used for visualization. Blue points indicate differential gene expressed in unexpanded cells, while red points indicate genes upregulated in expanded cells **E)** Differential gene expression in expanded (GZMB^hi^) and unexpanded (GZMK^hi^) CD8^+^ T cell subsets between Balinese and Papuan donors. Asterisks denote differentially expressed genes, while without are marker genes of cell subsets. Differential expression testing was performed using pseudobulk datasets with limma/voom.

Of the five level 2 cell types above, effector/memory CD8^+^ T cells were the only one with sufficient numbers of both expanded and unexpanded cells to allow direct comparison of expression profiles between the two expansion levels. We identified 416 significantly upregulated genes in expanded cells, and 401 in unexpanded cells at an *FDR* ≤ 0.05. These included a distinct set of marker genes previously associated with the different CD8^+^ subsets, validating our annotation strategy [61, 62]. *GZMB*, a molecule highly expressed in cytotoxic CD8^+^ T cells and associated with their cell-killing function was significantly up-regulated in expanded cells (*log*_2_*FC* = 0.52*, p* = 1.5 × 10*^−^*^9^). In contrast, unexpanded Effector/Memory CD8^+^ T cells exhibit upregulation of *GZMK* (*log*_2_*FC* = −1.00*, p* = 4.5 × 10*^−^*^14^), which has been associated with pro-inflammatory responses and activation of the complementary system (Figure 6D). Moreover, the proportion of GZMK^hi^ CD8^+^ T cells was significantly lower in donors from Denpasar and Sereh (Tukey’s post-hoc test *p <* 0.05), indicating population-level variation in this effector/memory subset (Figure S6). These findings demonstrate that TCR clone expansion is potentially useful in defining distinct subpopulations of effector/memory CD8^+^ T cells in peripheral blood in healthy individuals.

We then compared gene expression in both expanded and unexpanded effector/memory CD8^+^ T cells between the two provinces. In GZMK^hi^ T cells, interferon-stimulated genes (ISGs) including *IFITM1*, *IFITM2*, and *IFITM3* were up-regulated in Papuan donors relative to Balinese, while in GZMB^hi^ cells, *IFITM1*, *IFITM2*, *STAT1*, and *ISG20* showed higher expression in Papuan donors (Figure 6E). Notably, these ISGs are known to mediate the response to viral infection [63]. Other genes involved in viral-related pathways and Gene Ontology terms, such as regulation of viral genome replication and viral entry into host cell, included *DDX5*, *TNIP1*, and *STOM*, also had higher expression in Papuan donors (Table S14). These findings were consistent with gene co-expression network analysis, where a viral-related module in CD8^+^ T cells (Yoboi-1; Figure 5B-C) that includes *IFITM1* gene was observed to preserved only in Papuans.

### CDR3*β* sequences reveal local environment-associated TCR repertoire differences

Finally, we directly examined T-cell receptor sequences across individuals and sites, reasoning that different pathogen and lifestyle pressures across sites may have differentially impacted these. Specifically, we focused on the sequence of CDR3*β* in each T cell, as it determines antigen specificity of the TCR. Across all T cell subtypes the distribution of CDR3 sequence lengths in TCR *β* chains followed expected patterns, with most sequences ranging from 10 to 19 amino acids (Figure S5D). To assess the similarity of TCR repertoires across sites and individuals we calculated Jaccard indices based on the top 50 enriched motifs from each site separately for CD4^+^ and CD8^+^ T cells (Figure S7). Jaccard indexes for CD4^+^ T cell TCR repertoires were consistently 0 across all site pairs (Figure 7A), although limited sharing of CD4^+^ T cell TCR repertories is not unexpected. CD4^+^ T cells exhibit higher richness and lower clonality compared to CD8^+^ T cells, and they also possess a broader range of TCR repertoires [57, 64] Patterns of CD8^+^ T cell repertoires were more varied. The highest Jaccard similarity was observed between Sereh and Yoboi (0.024), followed by Denpasar and Pedawa and Denpasar and Sereh (0.011 in both cases). Similarity in cross-province comparisons was generally lower, with the lowest Jaccard indices between Yoboi and the groups Denpasar and Pedawa (0 in both cases). Similarly, Jaccard similarity within each site was highest among Denpasar donors compared to other sites, suggesting that the local environment shapes TCR repertoires (Figure 7A).

**Figure 7.**
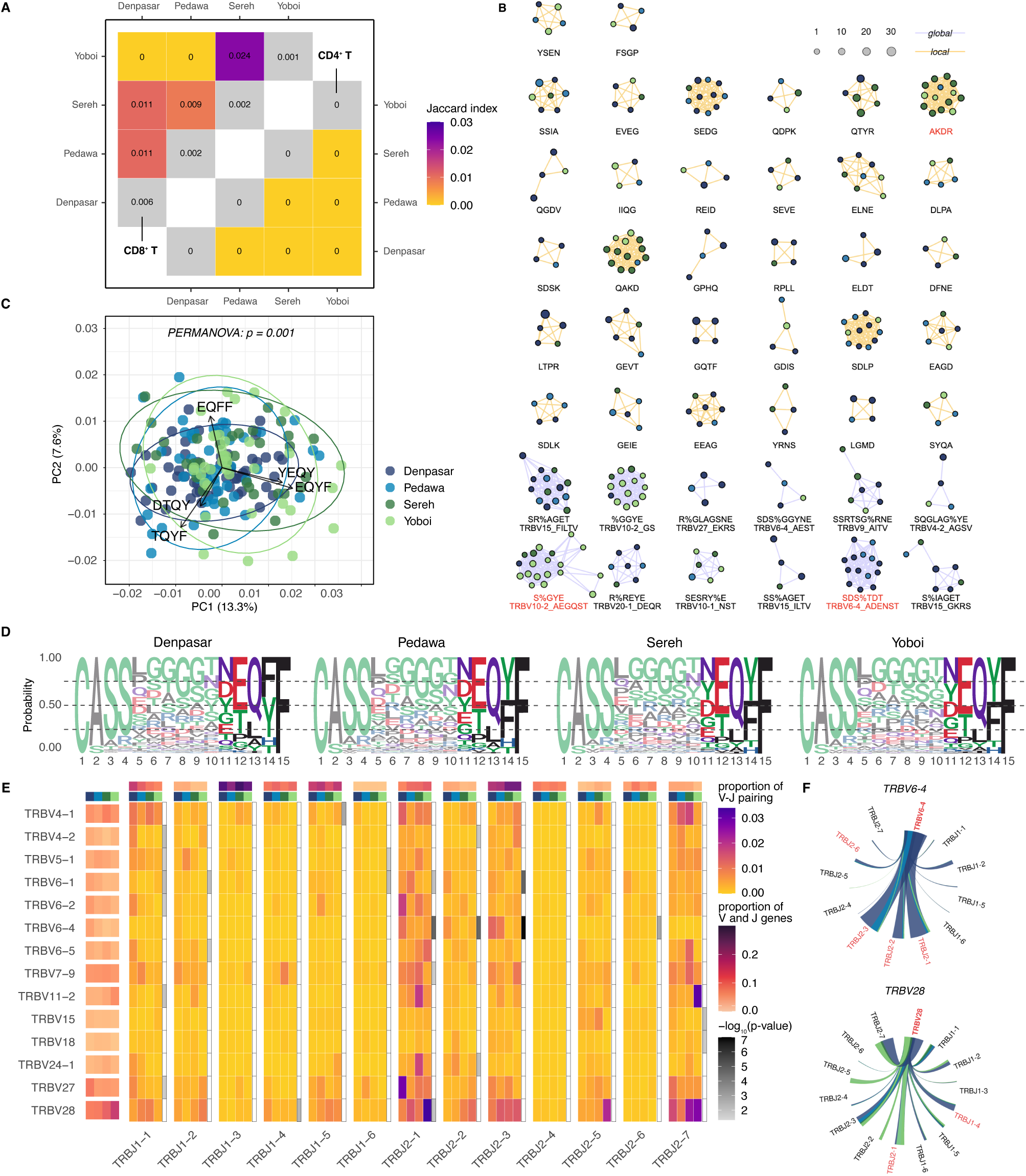
Differences in CD8^+^ T TCR repertoires between populations. **A**) Matrix of Jaccard similarities across top 50 enriched motifs in CD8^+^ and CD4^+^ T cells between sites. Diagonals represent mean Jaccard similarities of pairs of donors within each site. **B)** Global (purple) and local (orange) CDR3*β* sequence clusters. Only clusters containing at least four nodes are shown. Node colours indicate sampling site. Red indicates motifs with significant excess of individuals from one province. **C)** PCA of 4-mer composition from CDR3*β* sequences, showing the first two components. Arrows indicate the top five contributing loadings. Ellipses represent 95% confidence intervals around centroids of each site. **D)** Sequence logos of CDR3*β* region from all four sites. Amino acids are represented by single-letter codes and colour-coded by biochemical properties (red: acidic, blue: basic, black: hydrophobic, purple: neutral, and green: polar). **E)** Heatmap of significant V-J gene pairings. The y axis shows only V genes that are significant in V-J pairings, with proportions within each site. The x axis shows all J genes with proportions within each site. Grey scale represents adjusted p-values. **F)** V-J pairing examples at two different TRBV genes. Red indicates significantly skewed V-J pairings.

Building on these findings, we investigated CDR3*β* sequence sharing across all samples in these healthy cohorts. Cluster-based analyses with *GLIPH2* allowed us to identify site– and island-specific clonotypes and their potential biological relevance. We identified 32 clusters of locally similar—defined as sharing of an amino-acid 4mer regardless of position—CDR3 sequences and 12 clusters of globally similar—defined as sharing the entire sequence—sequences (Fisher’s exact *p <* 0.05) with at least four observations across the entire dataset; clonotypes grouped into a cluster are expected to share some antigen binding specificity (Figure 7B). We then examined the origin of individuals in each cluster. This analysis revealed that one local (AKDR, *p* = 0.005) and one global (S%GYE, *p* = 0.039) cluster were significantly overrepresented in Papuan individuals, while one global (SDS%TDT, *p* = 0.006) cluster was uniquely overrepresented amongst Balinese (all values from FDR-corrected hypergeometric test). We then searched for the two significant global clusters in VDJdb, a public curated database of TCRs with known antigen specificity [65]. The Papuan-enriched sequence S%GYE matched TCRs reactive to Cytomegalovirus (CMV), while the Balinese-enriched sequence SDS%TDT matched TCRs specific to Epstein-Barr virus (Table S15).

To further investigate sequence-level differences in full-length CDR3*β* regions, we performed PCA on the frequency matrix of all possible amino-acid 4-mers (including terminal ones) and examined clustering patterns across sampling sites. Although the cluster-based analyses above were generated after trimming of the N– and C-termini, PCA revealed that the top five sequences driving separation along PC1 and 2 were derived from the five C-terminal amino acids (Figure 7C). Notably, QFF and QYF motifs were dominant in urban and rural populations, respectively (Figure 7C-D). Consistently, PERMANOVA confirmed that PCA clustering by sites was statistically significant (*p* = 0.001).

We then analysed the pairing of V*β*-J*β* genes to assess whether differences in gene usage 18 TRBV genes and 2 TRBJ genes were more frequently observed in one island; for instance, *TRBV6-4* in Balinese and *TRBV28* in Papuans (Figure S8). In addition, 24 significant V*β*-J*β* gene pairs showed significant differences in pairing frequency across islands or sites (all FDR-corrected Kruskal-Wallis *p <* 0.05; Figure 7E). *TRBV6-4* was significantly paired with *TRBJ2-1*, *TRBJ2-2*, *TRBJ2-3*, and *TRBJ2-6*, while *TRBV28* showed preferential pairing with *TRBJ1-4* and *TRBJ2-1* (Figure 7E-F). Other V-J pairs that appear visually different but were not statistically significant, most likely because they represented rare occurrences with limited observations given our sample size. Nevertheless, our results suggest that environmental and genetic diversity across sites can directly impact V-J gene pairing and TCR diversity.

## Discussion

We have generated the first comprehensive single-cell resolution immune atlas from Indonesia, redressing the long-standing absence in human genomics of the world’s most genetically and environmentally diverse countries and regions. This publicly available resource captures regional genetic ancestry (East Asian-like and Papuan-like) and lifestyle diversity (urban and a multiplicity of rural settings), providing representation that is rarely, if ever, captured by existing datasets. Despite the challenges intrinsic to collecting and working with highly sensitive and labile samples outside urban Global North cities, our success in controlling for technical batch effect demonstrates that rigorous, cutting-edge functional genomics research in resource-limited settings is not only feasible but essential for understanding human immunity across the full spectrum of human diversity.

Our eQTL and colocalisation analyses revealed that a large majority of gene regulatory diversity is common between our cohort and other genetic ancestries. Nonetheless, we also identified a substantial number of eQTLs that did not colocalise across populations, despite using a relaxed threshold for colocalisation. While some of these are attributable to differences in sample size and cell type definitions across studies, they also included genes such as *ANKRD36B*, where signal in our cohort was driven by a set of eSNPs segregating at very low frequencies in individuals of European-like genetic ancestry. This example highlights the value of eQTL discovery in relatively small cohorts from diverse ancestries, akin to what has been observed in GWAS [66]. The importance of regional population structure was also evident in both our local ancestry analyses and our mapping of GxE interactions, where we consistently identified genes whose expression varied across individuals in an ancestry-dependent manner, many of them—including *MARCO*, *CDIP1*, *PLAAT3* or *SLC35A1*—directly involved in immune sensing. This signal is consistent with the hypothesis that population-differentiated allele frequencies at immune loci can reflect local adaptation or pathogen-driven selection, rather than neutral drift [67].

At deeper timescales, our results demonstrate that archaic hominins have also made significant contributions to immune gene regulation in ISEA. This includes robust Neanderthal signals at *IFIH1*, a gene where mutations often cause trade-offs between sensitivity to antigens and autoimmunity [68], and *TLR1*, a cell surface receptor that is key to the activation of the adaptive immune response and has been previously shown to be the target of adaptive introgression [49]. In parallel, and although introgressed Denisovan alleles can reach population allele frequencies in excess of 0.5, their complete absence in European-like genetic ancestry individuals means that their functional potential remains poorly understood. The inclusion of 87 individuals of primarily Papuan-like genetic ancestry in our study represents a 4-fold increase in the representation of individuals of this ancestry in RNA-seq datasets [6], allowing us to identify nearly 50 eGenes that are significantly associated with Denisovan ancestry tracts. The complex patterns of regulation and introgression that we identify around *GBP5*, *IL7R*, *HLA-E* or *STAT2* provide functional support to the theory that Denisovan introgression disproportionately contributed regulatory variation at immune loci in the Indonesian archipelago, especially in Papuan-ancestry populations, over tens of thousands of years of evolution [69, 52, 70]. All of these signals argue that both recent admixture and archaic gene flow have left significant traces on immune regulatory diversity across Island Southeast Asia.

Our results also point towards a consistent interplay between genetic and environmental forces in shaping steady-state expression levels in human immune cells. Although all participants in our study reported being healthy at the time of collection and showed no evidence of immune system activation, we nonetheless identified geographically-structured gene co-expression networks, as well as clear differences in TCR repertoires and clonal expansion patterns across sites. This suggests that the variable pathogen exposures and lifestyles that characterise human life in the Indonesian archipelago have readily quantifiable impacts on steady-state gene expression levels and TCR repertoire diversity, replicating previous observations on the existence of an immune state continuum across individuals living in urban and rural settings [20]. In our data this was most strongly evidenced by differences in steady-state expression levels of interferon-associated genes such as *IFITM1*, *IFITM2* or *IFITM3*, marked preferences in V-J gene pairings across sites and significant differences in CDR3*β* repertoire composition. Epitope matching revealed only associations with cytomegalovirus and Epstein-Barr virus in Papua and Bali provinces respectively, although this likely reflects the incomplete nature of epitope databases. While population structure drove some of the gene-by-environment signals that we observe, patterns of expression of other genes at the forefront of the host-pathogen interface such as *ERAP2* also suggest significant contribution to steady-state gene expression levels by forces beyond direct genetic effects. Overall, our results argue that evolutionary change operating at the DNA sequence level across large timescales shapes immune function differences but also highlight the responsiveness and short-term plasticity of the immune system that depends on local infectious pressure.

The sample size of our study is relatively modest compared to other recent large atlases [e.g. 19, 31, 71, 72], limiting our ability to detect subtler genetic effects or to examine rare cell types or states. Despite this we were able to identify unique regional signals that reflect Indonesia’s, and Island Southeast Asia’s lengthy and rich history of human habitation. More broadly, aspects of our work highlight the challenges of carrying out functional genomics research in resource-limited settings outside the Global North, and we faced substantial logistical barriers in its execution. For example, reagent costs were approximately twice as high in Indonesia as in Australia. Limited stocks, extended lead times, currency exchange fees and uncertainty in reagent and equipment access all contributed additional friction. These challenges are not unique to Indonesia: they are systemic barriers that limit research across the Global South. Realising the vision—and the potential—of functional genomics and genomic medicine will require concerted global commitment to workforce training, capacity building and infrastructure maintenance worldwide.

## Materials and Methods

### Ethics approvals, study design, and sample collection

This study was intended to profile immune cell differences representing variation across genetic ancestry and the environment in Indonesia. The study was reviewed and approved by the Mochtar Riady Institute for Nanotechnology Ethics Committee (#009/MRIN-EC/ECL/VI/2022), approval for data analysis in Australia was additionally granted by the University of Melbourne’s Human Research Ethics Committee (Approval ID 25179). We implemented a structured community engagement strategy to ensure that sample collection across all sites was conducted ethically and transparently. Prior to sample collection, we engaged with local stakeholders, including academic partners and archaeologists at Udayana University in Bali and Cenderawasih University in Jayapura, as well as with village and community leaders, local health officers, and hospital groups. Community briefings were held to explain the study objectives and sampling procedure in accessible language, with translation support from local partners when needed, and initial informed consent was obtained at the community level prior to individual participant recruitment. Individual-level informed consent for the collection and use of biological samples was additionally obtained from each participant prior to sample collection. We worked closely with health officers from the local Puskesmas (Community Health Center, part of Indonesia’s public healthcare system) and local hospitals to coordinate and conduct blood collection, and individuals were offered health examinations (finger-prick blood collection for measurement of blood biomarkers and screening for malaria and dengue) regardless of whether they chose to donate a blood sample to the study or not. 203 individuals agreed to participate in the study. From each of them we collected 10mL of whole blood in sodium heparin tubes (for PBMC processing) and 3mL in EDTA tube (for genomic DNA processing; both BD Vacutainers), as well as detailed phenotypic and lifestyle questionnaires. Blood samples were kept on ice until transferred to the lab for processing.

### PBMC isolation

PBMC isolation for Balinese samples took place at the Faculty of Medicine, Udayana University in Denpasar, Bali, with most samples (n=91) processed within 10 hours, and the rest (n=21) within 21–27 hours (Table S2). Due to limited access to laboratory facilities at Cenderawasih University, all samples were flown to MRIN in Jakarta (3,804 km away, approximately 6 hours direct flight time) no more than 24 hours after collection, such that times to isolation for the Papuan samples ranged from 13.2–38.5 hours (Table S2). Samples from the two provinces were processed by the same team members.

Once in the laboratory, blood samples were kept upright at room temperature until processed. We used fetal bovine serum (FBS; 10010023, Thermo Fisher Scientific) for the PBMC isolation, pooling, and washing procedures. Resuspension of cell pellets was done using a P1000 micropipette (Eppendorf, Germany). A total of 10 mL of whole blood was diluted 1:2 in phosphate-buffered saline containing 2% FBS (PBS + 2% FBS), followed by gentle inversion 4-5 times. The solution was added to a SepMate™-50 tube (Cat. No. 15450, STEMCELL Technologies, Vancouver, Canada) containing 15mL of Lymphoprep™ (Cat No. 7851, Stemcell Technologies), then immediately centrifuged at 1200 x g for 10 minutes at room temperature. Centrifugation was performed at 1200g for 20 minutes for selected blood samples collected overnight from Bali and for all samples from Papua, due to the presence of haemolysed plasma. The top layer containing PBMCs was collected and washed twice with PBS + 2% (300 x g, 8 minutes, room temperature). For cryopreservation, 4mL of cryopreservation medium (90% FBS + 10%DMSO) was added to resuspend the PBMC pellets, and the mixture was then divided into 4 x 1mL cryovials. The cryovials were transfered to liquid nitrogen for long-term storage.

### DNA isolation, genotyping, and genotype imputation

Genomic DNA from all samples was extracted using Gentra Puregene BloodKit (Qiagen,USA) following the manufacturer’s protocol. At least 100 ng of DNA per sample were sent to Gencove (NY, USA) for low-coverage sequencing. Raw FASTQ sequence reads were aligned to the GRCh38 human reference genome by Gencove, and the resulting BAM files were processed using standard quality control measures. Four of the samples from Sereh were sequenced with higher coverage to serve as imputation control. Sequencing libraries of those 4 samples were prepared using Illumina DNA Prep PCR-Free. 150bp paired-end sequencing runs were performed on the Illumina NovaSeq X with an average of 600M reads per sample with the expected mean depth of 20x. The alignment was performed using bwa-mem2 [73] using GRCh38 with alternative sequences, plus decoys and HLA for the alignment. All subsequent processes, including QCs, mark duplications, base recalibrations, haplotype– and genotype-calling were performed using GATK v4.5 best practice.

To maximize representation of relevant haplotypes, we constructed a composite imputation reference panel by merging and phasing variant data from three sources: i) 1000 genomes [74], ii) Indonesian genomes [3, 6], iii) Papua New Guinean genomes [10], totaling 1,464 individuals in the panel. All reference datasets were harmonized to the same GRCh38 build. Each chromosome of the merged reference panel was separately phased using Beagle v5.4 [75]. We then performed imputation using GLIMPSE2 [76], following recommended best practice. Chromosomes were divided into imputation chunks using GLIMPSE2_chunk_static. Genotype likelihoods (GLs) were computed for all target samples at all variant sites present in the reference panel, and only autosomal bi-allelic SNPs were considered. Imputation and phasing were performed using GLIMPSE2_phase_static for each sample and chromosome chunk. After per-chunk imputation, output BCF files for each chromosome were concatenated using GLIMPSE2_ligate_static to produce a single, phased, and imputed BCF file per chromosome per sample. The final output consisted of autosome-wide, phased and imputed BCF files for each sample. Imputation accuracy was assessed using GLIMPSE_concordance by comparing selected imputed genotypes to available high-coverage data from the same individuals.

### Local ancestry inference and archaic introgression calling

Genetic ancestry in Indonesia is broadly characterised by an longitudinal ancestry cline that transitions from “East Asian-like” genetic ancestry in the west to “Papuan-like” genetic ancestry in the east as a consequence of the region’s demographic history [77] and depth of human habitation. Thus, we performed genome-wide local ancestry mapping for all samples using reference haplotypes drawn from populations outside the study samples. For the East Asian-like reference, we combined 1000 Genomes CHS, CDX, and KHV [74] together with Indonesian Mentawai samples, which were genetically close to the Austronesian-related proxy groups [3]. The Papuan-like reference was a composite of highland Papuan populations [10]. Reference haplotypes were kept separate from the study samples. We ran RFMix v2 [22], a discriminative approach that models ancestry along an admixed chromosome given observed haplotype sequences of known ancestry, on phased haplotypes in a population-aware mode by treating the two reference groups as training populations and the study samples as unknowns. We set the terminal node size with the option –n 5 to control the random forest node, and all other RFMix parameters were left at their defaults.

Archaic introgression was inferred from imputed genomic data using a Hidden Markov Model based method implemented in HMMix [43]. HMMix finds regions of elevated variant density remaining after removing the variation found in the putatively non-introgressed outgroup [43]. The tool was run for each individual with the –phased option using provided archaic and outgroup variants [78]. Inferred archaic genomic segments were filtered by the probability of incomplete lineage sorting (ILS) as previously described [79, 50], which led to the removal of 35% of identified segments (14% of total length). Recombination rate for each archaic segment was inferred from the deCODE recombination map [80] by weighted average. The impact of the recombination map choice was assessed by using the maps from another study [50]. Changing the recombination map altered the outcome of filtering in 10% of cases on average. The time of population split between ancient modern humans and the common ancestor of Denisovans and Neanderthals was set to 550kya as the most recent estimate that is covered by multiple different approaches for more stringent filtering [81, 82, 83, 84, 85]. We used 70kya for the age of introgression as the most ancient estimate [81, 3] which still falls within the dating range for Denisova 3 remains [86]. Significance was corrected by FDR. To evaluate the performance of introgression calling in low-pass imputed modern human genomes, we compared the introgressed segments inferred from low-coverage and high-coverage data in 4 Sereh individuals for whom high-coverage genomes were available (Figure S4A,B). The amount of overlap between low coverage and high coverage data was calculated with bedtools intersect [87] (Figure S4A). Fragments with fewer than 10 SNPs were excluded from further analysis.

The ancestry of each archaic fragment was assigned based on the predominant number of Denisovan [88] or Neanderthal alleles in each introgressed segment. The number of Neanderthal alleles was calculated as the maximum from three high-coverage Neanderthal genomes from Vindija, Chagyrskaya, and Denisova caves [81, 89, 82]. However, for the contour density plots (Figure S4C), the average from three Neanderthal genomes was used since otherwise, the plot collapses in the absence of Denisovan introgression in Balinese individuals. If the number of matching archaic variants was less than 5, the ancestry was assigned to “unknown” Plotting the proportion of SNPs matching Neanderthal or Denisovan genomes as normalised density shows that the assigned ancestry of the segments is generally well-resolved (Figure S4C).

### Population demographic inference

We computed the pairwise kinship matrix using KING v.2.3.1 [90]. For demographic analyses, we removed an individual from pairs that exhibit close relationships (less than 2nd degree). We used ADMIXTURE v1.3 [91] to observe population structure by classifying individuals into K clusters based on genetic similarity using maximized likelihood with high-dimensional optimization. Ten randomly seeded runs were performed for each number of ancestral populations (K = 2–15), and the results within each K were summarized with CLUMPP v1.1.2. Cluster K = 8 was shown to have the lowest cross-validation error value and was therefore chosen as the best model describing the dataset. To formally estimate the proportion and the time of admixture, we performed F4Ratio [21] and ALDER [92] analyses respectively. To estimate the population size dynamics, we conducted MSMC2 [93] analysis. Based on MSMC2’s cross-coalescence results, we then performed the MSMC-IM [23] analyses to infer split times between a pair of populations. It fits a time-dependent migration model to the pairwise rate of coalescences based on estimates of coalescence rates within and across populations.

### Single-cell library construction and sequencing

We generated single-cell RNA-sequencing (scRNA-seq) data using the Next GEM Single Cell 5’ Kit v2 and Next GEM Chip K from 10X Genomics. TCR amplification libraries were made in conjunction with the with the Next GEM Single Cell 5’ Kits in accordance with 10X Genomics protocol CG000331 Rev F. Each sequencing library contained 8 different PBMC samples. Cryopreserved PBMCs were thawed in batches following Demuxlet Cell Preparation Protocol V.2 [94] and viability was assessed using a Countess II(Cat. No. AMQAX1000, Thermo Fisher Scientific). For each batch we aimed to pool 240,000 cells from each individual into a single aliquot; however, if any individual had fewer than 240,000 cells, we matched this amount across the remaining samples in the batch to maintain equal representation of donors. Batches were balanced across sampling site, age and sex as much as possible, with most batches containing two randomly chosen samples from each site and no batch containing samples from only one site. Commercial human PBMCs from Lonza (Lonza 4W-270, lot #3038099) added as controls at a 1:4 ratio to each main sample (e.g. when pooling 240,000 cells per sample across 8 samples, 60,000 Lonza control cells were added. This control was added for potential integrated analyses with other atlases using the same control, such as the Asian Immune Diversity Atlas (AIDA) Data Freeze v2 dataset [19]. From each library’s pool, we loaded up to 70,000 cells into a well on the chip. For samples or libraries with <70% average cell viability, we either repeated the entire low-quality library or merged low-viability samples into new multiplexed libraries. Overall, we generated 29 sequencing libraries across 203 Indonesian donors. All libraries (both 5’ and TCR) were run on NovaSeqX using 300 cycle 10B flow cells, with the read parameters 151:11:11:151.

### Single-cell RNA-sequencing (scRNA-seq) data pre-processing and quality control

Raw paired-end FASTQ files from each 5’ scRNA-seq library were processed using 10X Genomics Cell Ranger version 7.2.0 with introns included in the cellranger count step. Reads were mapped to Cell Ranger human reference 3.0.0 (GRCh38, Ensembl v93, date 2018-11-19). Based on the number of cells estimated and average viability during GEM library preparation, we removed 2 sequencing libraries and only used 27 libraries for further analyses, still representing all 203 donors.

We used *Demuxafy* [95] for genetic demultiplexing and droplet assignment. We used vireo [96] for genetic demultiplexing at the BAM and barcode level from each sequencing library’s filtered Cell Ranger output in two different ways. To identify cells from the commercial Lonza PBMC control, we first ran vireo on genotype-free mode and assigned cells based solely on common SNPs with *M AF >* 0.05 using the 1000 Genomes Phase 3 reference file, as recommended by *Demuxafy* [95], then donor-matched the predicted clusters with array-based genotype data of the exact lot number (Lonza 4W-270, lot #3038099) generated by the Asian Immune Diversity Atlas (AIDA) for their Phase 1 data [19]. We then ran vireo a second time on paired-genotype mode and assigned cells to all remaining clusters based on the paired WGS data. For droplet assignment, we used scds [97], scDblFinder [98] and DoubletDetection [99], from which we merged the results with vireo and inferred majority singlet assignment. At this stage we filtered individual cells to retain only those with at least 3,000 detect genes (*NODG*) ≥ 300 and percentage of mitochondrial reads (*pMito*) ≤ 10%. We then performed preliminary cell type annotation using RCAv2 [100], which projects each library to immune cell reference panels, as demonstrated by AIDA [19]. Each sequencing library was processed as a separate Seurat v5 [101] object prior to integration.

### Data integration and cell type annotation

Integrative analysis was done using Seurat v5 with Harmony version 1.2.0 [102] as the integration method. We implemented a sketch-based integration approach [103], subsampling 5,000 cells per sequencing library, to allow for a more resource-efficient integration while still preserving the representation of both abundant and rare cell types in each library. This approach used BPCells [104] to manage memory requirements. At this point we removed cells that had been assigned to Lonza PBMCs prior to further analyses, as well as four Indonesian samples with *<* 500 cells, leaving a total of 199 individuals for all downstream analyses. To characterize the cells more accurately, we performed manual cell type annotation using canonical marker genes (Table S3) based on different Seurat clustering (resolutions: 0.1, 0.2, 0.3, 0.5, 0.7, and 1). Annotation was done at three different levels: manual cell annotation on all cells into broadly defined cell types, supported by Seurat clustering at 0.3 resolution (level 1), more detailed lineage-specific lymphoid-only and myeloid-only annotations (level 2), and a deeper annotation on lymphoid cells using markers from a previous rural-urban study and immune cell reference panels by AIDA cohort (level 3; [20, 19]).

### Cell type proportion analyses

We used propeller [105] to calculate transformed proportions of each level 1 cell type per sample, and correlated proportions of each cell type with known biological (e.g. age, sex, BMI) and technical (e.g. sequencing batch and time to PBMC isolation) covariates. To assess significant cell type-specific differences between islands and sampling sites, we used propeller’s moderated t-test and included age, sex, sequencing batch, and time to PBMC isolation as covariates. We also assessed whether there are significant cell type-specific differences between sex, age, and BMI after controlling for other covariates. For visualization, we regressed out the covariates from the transformed proportions and plotted the residuals.

### Cell type-specific differential gene expression analysis and adjusting for unwanted technical variation

To account for the possible confounding effect of variation in the time to PBMC isolation between sampling sites we used *RUVg* [25]. We pseudobulked count data to aggregate per donor for each cell type, keeping only genes with *log*_2_*CPM >* 1 and expressed in at least 50% of donors in at least one sampling site. We used the list of single-cell stably expressed genes (SEGs) included in the scMerge package [26] as controls. The recommendations for using RUVg require determining an ideal number of *W* s to include in downstream analyses; this value should be calibrated to control for unwanted variation while preserving variation of interest in the dataset. To determine a suitable *W* value for our dataset we took advantage of the fact that the times to isolation for samples from Pedawa and Sereh were bimodally distributed (Figure 2A). This was most extreme in Pedawa, where 32 samples were processed within 10 hours of collection (“early”, median = 6.55 hours) and 20 samples after 10 hours (“late”, median = 23.65 hours). In Sereh, 17 samples were processed within 20 hours (“early”, median = 16.8 hours) and 25 samples after 20 hours (“late”, median = 29.5 hours). As all donors reported being healthy at the time of collection, we expect few biological differences between samples from the same site that were processed early or late. We performed differential expression testing between early and late samples from Pedawa and Sereh using a model that included age, sex, sequencing batch, cell count, and time to isolation as covariates in limma/voom version 3.62.2 [106, 107], and used TMM normalization as implemented in edgeR version 4.4.2 [108].

We considered values for *W* from 6 to 10. For each value of *W* we repeated differential expression testing using a model that incorporated age, sex, sequencing batch, cell count, time to isolation and all unwanted variation factors as additional covariates, and compared these results to those from the model without the W factors. *W* = 8 as it struck a balance between removing the most unwanted technical variations while preserving the biological differences between sampling sites (Figure S2A). All differential expression testing between the four sites or between cell types was performed using limma/voom with TMM normalisation and the full model including the 8 *W* factors. Genes with an FDR-adjusted *p* − *value <* 0.05 were considered differentially expressed.

### Expression quantitative trait loci (eQTL) mapping and sharing

We used quasar [28] v1.0.0 to identify eQTLs in nine level 1 cell types with log_10_(median cell count) *>* 1.3, which included CD4^+^ T, CD8^+^ T, NK, CD14^+^ monocyte, memory B, naïve CD8^+^ T, naïve B, CD16^+^ monocyte, and DC. Genes expressed in at least 1% of all cells, and in 10% of individuals in a given cell type were considered for testing. We excluded SNPs with a minor allele frequency (MAF) <0.05 across the entire dataset, genotype missingness >10%, or those with Hardy-Weinberg Equilibrium (HWE) *p <* 10*^−^*^5^. Due to the observed levels of cryptic relatedness and high levels of population structure in our cohort (Table S4), we used a linear mixed model approach and included the kinship matrix as a random effect. We used log-normalized mean-aggregated pseudobulk for each sample per cell type, as recommended by the quasar documentation when using an LMM. Our final eQTL testing model also included age, sex, 10 genotype PCs, and 8 RUVg (*Ws*) as fixed effects. Quasar combines all SNP-level tests within a single gene using ACAT [29] to calculate gene-level p-values. We used these values, after FDR correction, to identify genes with a significant eQTL in the dataset.

To improve effect-size estimation and mitigate variation in statistical power across cell types, we applied multivariate adaptive shrinkage (*mashr*; [30]) on our quasar results to jointly model shared and cell type-specific eQTL effects. Residual correlation across cell types was estimated from a random subset of up to 10,000 variant–gene pairs, and data-driven covariance structures were learned from strong associations (per-gene FDR < 0.05 based on the minimum p-value across cell types), together with canonical covariance matrices. The fitted model was applied to all variant–gene pairs to obtain posterior mean effect sizes, standard deviations, local false sign rates, and posterior probabilities of negative effects for each cell type. Based on the mashr results, we considered two definitions of signal sharing between provinces or cell types: lead eSNP significance (local false sign rate; LFSR) and lead eSNP effect size (beta). We defined sharing by whether the tested eGene is also significant in the other population. For effect size, we defined sharing by the ratio of beta within a factor of 0.5 when compared to the other population.

Finally, we used GEMMA [109] v0.95.8 to identify genes with significant genotype by environment (GxE) effects, as quasar cannot directly model genotype interactions. For each ACAT-significant eQTL, we fit an LMM with GEMMA testing for an interaction between island and genotype at the lead eSNP. This model contains exactly the same covariates as those used in the original quasar fit, including the kinship matrix, with the genotype-by-island interaction specified using the *-gxe* option. LRT-derived p-values from this model were then FDR corrected to identify genes with significant GxE interactions.

### Local ancestry and archaic assignments to eQTLs

To identify eQTLs driven by local ancestry or archaic hominin introgression across the complete dataset we used the set of top SNPs identified by mashr above. We scanned each SNP for association with local ancestry by correlating haploid allele presence and haploid local-ancestry calls from RFMix’s msp files. For each site we built two aligned matrices: genotype G (variants x haplotypes, 0/1 for ALT on each haplotype) and ancestry A (variants x haplotypes, 0/1 for Papuan ancestry). Row-wise Pearson’s correlations between the G and A were computed and summarized using the covariance formula 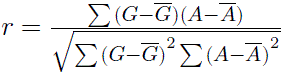. The squared correlation *r*^2^ is reported as the fraction of allele variance explained by Papuan ancestry. Per-variant p-values were calculated and multiple-testing corrected using the Benjamini-Hochberg procedure.

We also identified candidate archaic-introgressed eQTLs by testing whether the ALT allele is enriched on haplotypes inferred as Denisovan or Neanderthal by HMMix [43] above. Phased genotypes were converted into per-haplotype allele calls and intersected with per-sample HMMix introgression tracts. Each SNP and haplotype pair was annotated as introgressed or non-introgressed based on overlap with ancestry-specific HMM segments. For each site, we counted ALT and reference alleles on introgressed and non-introgressed haplotypes. Sites with insufficient data (e.g. very few introgressed haplotypes (introgressed haplotypes *<* 5) or ALT observations (ALT in introgressed haplotypes *<* 2)) were removed by independent filtering to retain only tests with power. We also removed alleles that segregated at frequencies ≥ 0.005 within the sub-Saharan African populations in the 1000 Genomes Project. We evaluated enrichment using Fisher’s exact test on the 2×2 table of allele counts and corrected p-values using the Benjamini-Hochberg procedure. Candidate introgressed SNPs were defined using thresholds combining statistical significance (*FDR >* 0.05), high ALT frequency on introgressed haplotypes, low ALT frequency on non-introgressed haplotypes, and consistency across populations.

### Colocalisation and fine mapping

To assess signal sharing between eQTLs identified across our entire sample and One1K1 [31], a cohort of 982 northern European genetic ancestry, we performed colocalisation and fine-mapping analyses using *coloc* v5.3 [27, 110]. For this, we downloaded OneK1K summary statistics from the eQTL Catalog (dataset IDs QTD000606-QTD000629, [111]). Not all OneK1K cell types had equivalents in our dataset; thus we only considered 14 cell types with equivalence to our level 1 annotation, detailed in Table S6. We considered all genes with significant eQTLs in Indonesia as defined by *FDR* − *correctedACAT p <* 0.05; of these, summary statistics from OneK1K were available for 76% of eGenes per cell type on average. We used the coloc.abf function with default settings, including setting the prior probability that a SNP was a significant eQTL in either dataset being tested to 10*^−^*^4^. We note that sometimes no SNP-level quasar p-value reached this threshold although the gene was significant according to ACAT; we excluded these genes from further analyses. We did not filter genes where the OneK1K summary statistics failed to reach this threshold.

|coloc.abf| does not make explicit use of linkage disequilibrium information, and assumes there is only a single causal variant in each region. Given our limited sample size and the fact that our genotype data was imputed to high coverage from low-pass sequencing data we elected to use this approach over the more sophisticated SuSiE [112], even though this would have allowed us to identify multiple causal variants with more confidence. Coloc outputs five posterior probabilities, *H*0, *H*1, *H*2, *H*3 and *H*4. These indicate the posterior probability that there is no causal variant in the interval being tested (*H*0), that there is signal in the first (*H*1) or in the second (*H*2) dataset only, that there are different signals in both populations (*H*3) or that there is a single shared causal variant between the two datasets being compared (*H*4). We considered genes where *H*4 ≥ 0.5 to be successfully colocalised and share a single causal variant, and genes where *H*1 or *H*3 *>* 0.5 as failing to colocalise and thus candidates for harbouring population-specific signals. Genes where no posterior exceeded 0.5 were not considered further. Fine-mapping was performed in the Indonesian dataset only using the coloc coloc.abf function with default settings. For each gene, we identified the top SNPs by posterior probability, as well as 95% PIP sets. As above, we excluded genes where no single SNP had a quasar p-value *<* 10*^−^*^4^, as many of these resulted in 95% sets that included 100s or 1000s of SNPs.

### Cell type-specific gene co-expression network analysis

We constructed gene co-expression network from the pseudobulked dataset after RUVg using the high-dimensional Weighted Gene Co-expression Network Analysis (*hdWGCNA*) package version 0.4.05 [113]. The weighted genes co-expression network framework constructs a gene-gene correlation matrix and generates a scale-free network based on the selected soft-thresholding power [114]. Network or a module is an undirected graph in which nodes represent genes and edges represent the gene-gene association. The hdWGCNA R package builds on Seurat’s data structure and functionality while maintaining full compatibility with Seurat. Co-expression networks were constructed separately for CD4^+^ T, CD8^+^ T, NK cells, Memory B cells, and Naive B cells across the four sampling sites. We excluded donor DPS-006 because its predominantly formed inflamed clusters, which could introduce donor-specific bias in the co-expression analysis.

We omitted the metacell construction step and instead generated a matrix of expression residuals for each site-cell type combination. For each combination, the soft-threshold power was calculated using the function TestSoftPowers and set to the value at which the scale-free topology index (R^2^) reached 0.8. To reduce the number of modules containing fewer than five genes, we independently adjusted the minimum module size (ranging between 20-50) and deepSplit (ranging between 2-4) parameters for each site. Specifically, we used a MinModuleSize value of 50 and a deepSplit value of 4 for Denpasar, 20 and 4 for Sereh and Pedawa, and 50 and 2 for Yoboi. To identify modules that were shared or not shared in other groups, we tested module preservation using ModulePreservationNetRep [115]. We calculated an overall preservation score by averaging the seven attribute p-values, adjusting for FDR, and classified modules with an *FDR* ≤ 0.05 as preserved. Each module was annotated using EnrichR, with GO Biological Process (2025) gene sets to characterize biological processes and MSigDB Hallmark (2020) gene sets to assess functional pathways. Network plots were constructed using all genes in each module but visualised by displaying only gene–gene connections with edge weights > 0.4. Hub genes for each module of interest were identified by selecting genes with *kME* ≥ 0.8 and were visualized in the center of network plot. In the network plot, each node (gene) was colored according to whether it was differentially expressed relative to other groups.

### Single-cell V(D)J-sequencing (scV(D)J-seq) data pre-processing and quality control

For each TCR-seq output, paired-end FASTQ files were processed using Cell Ranger VDJ pipeline version 7.2.0 and mapped to V(D)J reference version 7.1.0 (vdj_GRCh38_alts_ensembl-7.1.0). Pre-processed V(D)J data was combined with scRNA-seq cell barcodes to assign T cell receptor (TCR) sequences back to cells and individuals. We excluded TCR-containing cells with low-quality TCR sequences, including those with fewer than 2,000 TCR reads per cell, those where the CDR3 region did not start with cysteine and end with either phenylalanine or tryptophan [116], and orphan T cells possessing only one TCR chain or more than one pair of TCRs using *scRepertoire* package version 2.2.1 [117]. Additionally, we excluded DPS-006 from all TCR analyses because the majority of its lymphoid population consisted of inflamed cells, which could bias TCR analyses in our otherwise self-reported healthy cohort.

### T cell receptor (TCR) repertoires diversity analysis

We analysed TCR repertoires for both similarity and diversity using *immunarch* package version 0.9.1 [118] separately in each cell type. Clonal similarity analyses were conducted at cell type annotation level 1 in both CD8^+^ and CD4^+^ T cells. Overlap in clonotypes between individuals was quantified using the Jaccard index, and mean values were calculated to represent clonal similarity within each site. Clonal TCR diversity analyses were performed at cell type annotation level 1 and 2. TCR diversity accounts for both the number of clonotypes, referred to as richness, and the distribution of their abundances, referred to as evenness. To capture these components we employed three complementary indices. Chao1 is an estimator that quantifies clonal richness by estimating the total number of clonotypes. The inverse Simpson index integrates both richness and evenness, representing the effective number of equally abundant clonotypes based on their relative abundances [119, 120]. Gini coefficient measures the inequality in clonotype frequencies, serving as an indicator of clonal distribution within donors [119, 121].

We examined the effects of sex, age, BMI, and sampling site on diversity of TCR were then examined for their associations with diversity indices across all cell types. To account for potential confounding, all indices were calculated after regressing out covariates prior to analysis: sex was adjusted for age, sampling site, and time to isolation; age was adjusted for sex, sampling site, and time to isolation; BMI was adjusted for sex, age, sampling site, and time to isolation; and sampling site was adjusted for sex, age, and time to isolation. Residuals were used for visualisation and for testing pairwise differences using one-way ANOVA, followed by Tukey’s post-hoc test.

### Integration of scV(D)J with transcriptomic profiles

To investigate the relationship between clonal expansion and gene expression profiles, scRNA-seq data were integrated with TCR-seq using the *scRepertoire* package. This integration enabled us to annotate specific cell types based on gene markers from clonal expansion status (level 3 annotation). Clonotypes were classified as expanded when their frequency exceeded 1.5 times the median clonotype frequency per sample, following [122].

Among all T cell subsets, effector/memory CD8^+^ T cells were the only cell type with an average ≥ 10 cells per individual in both expanded and unexpanded cells. This was sufficient to allow fair comparison of gene expression between two cell types. Differential expression between expanded and unexpanded effector/memory CD8^+^ T cells was assessed using a model that included age, sex, cell count, time to isolation, and 8 *W* factors as covariates in limma/voom as described above. Significantly differentially expressed genes between expanded and unexpanded effector/memory CD8^+^ T cells were used to annotate GZMK^hi^ and GZMB^hi^ CD8^+^ subsets. Furthermore, we analysed DGE in both GZMK^hi^ and GZMB^hi^ CD8^+^ T subsets between two provinces. Significantly differentially expressed genes were subjected to Gene Ontology (GO) enrichment analysis using *clusterProfiler* package version 4.14.6 [123]. FDR correction was then applied to adjust for multiple testing in enriched pathways [124].

### Complementarity-determining region 3 sequence composition and motif enrichment analysis

To investigate TCR specificity between populations, we focused on the CDR3*β* chain in both CD8^+^ and CD4^+^ T cells. Motif enrichment analysis was performed within each sampling site independently using *HetzDra/turboGliph* version 0.99.2 [125, 126]. We then calculated Jaccard similarity among the top 50 enriched motifs based on Fisher scores between sites, and analysed enriched motifs between sites to identify population-specific clusters in CD8^+^ T cells.

We used *GLIPH2* to identify clusters based on sharing of global (the complete CDR3*β* sequence after excluding the first and last three amino-acids) or local (amino-acid 4-mers in any position after excluding the first and last three amino-acids) motifs using the following parameters: simulation depth = 1000, k-mer minimum depth = 3, and the default *GLIPH* reference file. Clusters were filtered using Fisher’s exact test with a significance threshold of *p <* 0.05. Population-specific clusters were assessed using a hypergeometric test based on the number of samples within each cluster, and corrected for multiple testing (FDR 5%). For epitope annotation of significant population-specific global clusters we input trimmed CDR3*β* sequences into VDJdb [65] with wildcard position (%) expanded to all possible amino acids. Full-length CDR3*β* sequences was included to ensure accurate mapping.

All local motifs identified by *GLIPH2* were 4-mers. To further investigate 4-mer patterns, we generated count matrices of all CDR3*β* 4-mers in the dataset using *immunarch*. Samples containing fewer than 30 CD8^+^ T cells were excluded due to low coverage, leaving 182 samples for downstream analysis. Empty values within individuals were imputed with the mean frequency of that particular 4-mer to enable PCA analysis on a complete matrix without loss of 4-mer components. K-mer counts were normalised to proportions per sample, and PCA was performed using the *prcomp* function in R to explore sequence composition across sites. PERMANOVA was applied on PC1 to PC10 using *vegan* package version 2.7-1 [127] to assess whether principal components were associated by sites. Sequence logos for each site were generated based on the most abundant sequence length to represent motif composition using *ggseqlogo* package version 0.2. Lastly, we analysed the frequency of specific V-J gene pairings within each individual. Significant V-J pairs across four sites were identified using a Kruskal-Wallis test, with FDR correction for multiple testing. V genes involved in these significant V-J pairs were then visualised in a heatmap. Circos plots of significant plots generated with the *circlize* package version 0.4.16.

### Data and code access

1x WGS, scRNA and TCR sequencing reads are currently being submitted to the European Genome-Phenoe Archive (EGA; accession number is pending). Genome-wide quasar eQTL summary statistics are available at https://doi.org/10.26188/31302163. Processed expression data has been deposited in CELLXGENE and is freely available to browse at https://cellxgene.cziscience.com/collections/d1e0e64d-6d2a-4a3e-b7f4-43ed909a9d9c. Analysis code for the results presented in this work is available in https://gitlab.svi.edu.au/igr-lab/indo_introgression (archaic introgression inference), https://gitlab.svi.edu.au/igr-lab/Indo_sc_expression (single-cell gene expression QC and initial downstream analysis), https://gitlab.svi.edu.au/igr-lab/Indo_sc_eQTL (eQTL calling and associated analyses), https://gitlab.svi.edu.au/igr-lab/indo_sc_vdj (single-cell TCR QC and downstream analyses).

## Acknowledgments

We are extremely thankful to all donors in Denpasar, Pedawa, Sereh and Yoboi, and in particular to the community elders, village officials, and organisers at Pedawa, Sereh and Yoboi, without whose participation this project would have been impossible. We thank Lisye Iriana Zebua, Hanny Felle, Terida Tokoro, Dr Aprilia Sokoy, Dr Ivone Rumasewu, Trias Pala, Anggraeni Ambarwati, and Kaleb Bonyadone for the assistance during community engagement and sampling in Yoboi and Sereh villages. We thank Jeffrey Pullin, Davis McCarthy and Jarny Choi for assistance with eQTL mapping and quasar, and members of the Gallego Romero group and the Genome Diversity and Diseases group at MRIN, as well as Shyam Prabhakar, Boxiang Liu, Kian Hong Kock, and additional members of the Asian Immune Diversity Atlas consortium for comments. This work was supported by Chan Zuckerberg Initiative award 2021-240137 to NB, MPC, SGM, HS and IGR; NHMRC Ideas grant 2020501 and ARC Discovery Project DP200101552 to IGR; Royal Society of New Zealand Marsden Grant 20-MAU-017 to MPC and IGR. PS is supported by a Commonwealth through an Australian Government Research Training Program Scholarship and St Vincent’s Institute Top-up Scholarship. PK was supported by Wellcome Trust International Training Fellowship 222992/Z/21/Z. St Vincent’s Institute acknowledges the infrastructure support it receives from the National Health and Medical Research Council Independent Research Institutes Infrastructure Support Program and from the Victorian Government through its Operational Infrastructure Support Program. The funders had no role in study design, data collection and analysis, decision to publish, or preparation of the manuscript.

## Author contributions

- Conceptualization: NB, IGR, MPC, HS, SGM, PK, CCD
- Formal analysis: MF, PS, PK, MMN, AG, IGR
- Funding Acquisition: NB, MPC, SGM, HS, IGR
- Data Curation: MF, PS, LP, RK, EM, LPe
- Investigation: IAl, IAp, CD, AC, MF, RK, PK, PCL, EM, MN, SO, LPe, LPr, BU, DMW, NNAD, AED, FS, SAKF, SGM, HS, IGR
- Project Administration: MF, MMN, HS, SGM, IGR
- Resources: IAl, IAp, CD, AC, MF, RK, PK, PCL, EM, MN, SO, LPe, LPr, BU, DMW, NNAD, AED, FS, SAKF, NB, SGM, HS, IGR
- Supervision: IGR, NB, SGM, HS, MPC, MF
- Visualization: MF, PS, PK, MMN, AG, IGR
- Writing – Original Draft Preparation: MF, PS, PK, MMN, AG, IGR
- Writing – Review & Editing: All

