## Supplementary figures 1-8 for "Ancestral and environmental diversity shape the immune landscape in Indonesia"

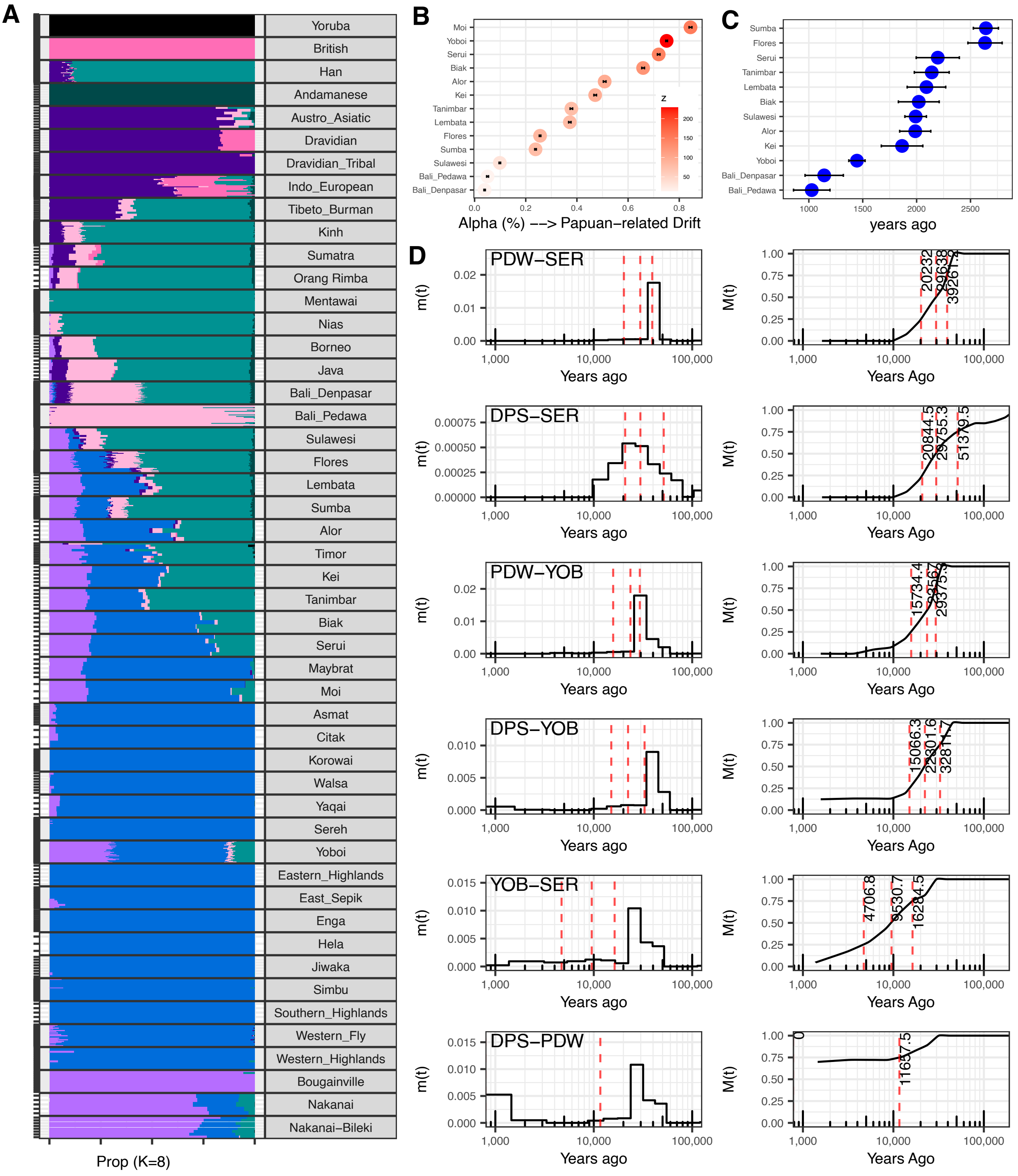


**Supplementary Figure 1.** Demographic analyses of the dataset. **A)** Population structure in our dataset as inferred using ADMIXTURE in K=8. **B)** Papuan-related ancestry proportion and **C)** time of admixture of East Asian- and Papuan-related ancestry in populations in our dataset, including populations in Eastern Indonesia and West Papua in the comparative datasets as inferred using F4ratio and ALDER respectively. **D)** Inference of population split using cross-coalescence analysis using MSMC-IM between population pairs in our dataset.

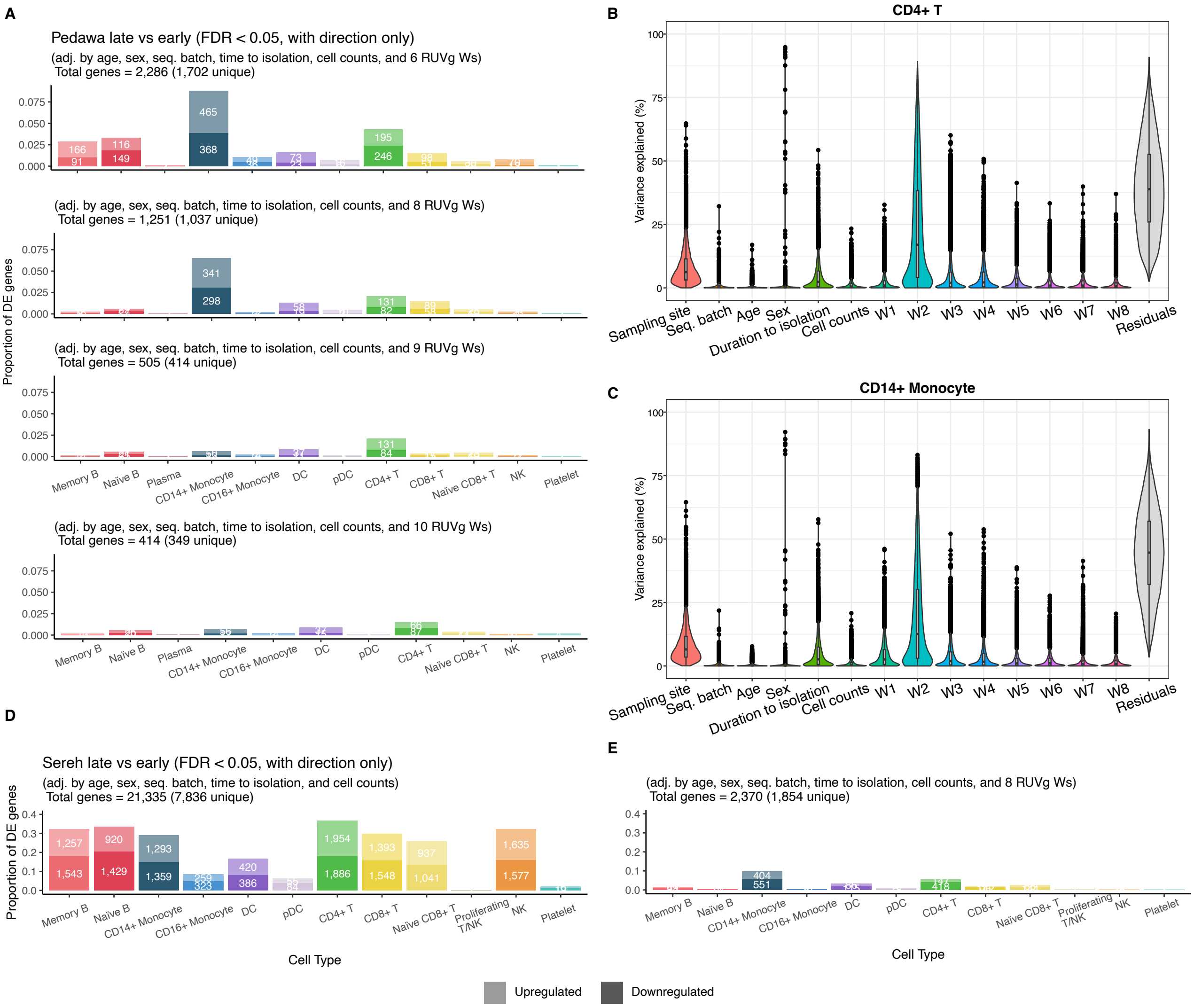


**Supplementary Figure 2.** RUVg results to assess and remove unwanted technical variations from the data. **A)** Comparison of the impact of different *k* values used as covariates in cell type-specific differential expression (DE) analysis between Pedawa late vs early donors. Out of the *Ws*, *W2* explained the most variance among all the covariates, as can be seen in **B)** CD4$^{+}$ T and **C)** CD14$^{+}$ Monocytes. **D-E)**Reduction in DE genes also found when comparing between Sereh early (<20h) and late (>20) (D) with no RUVg correction and (E) with RUVg correction.

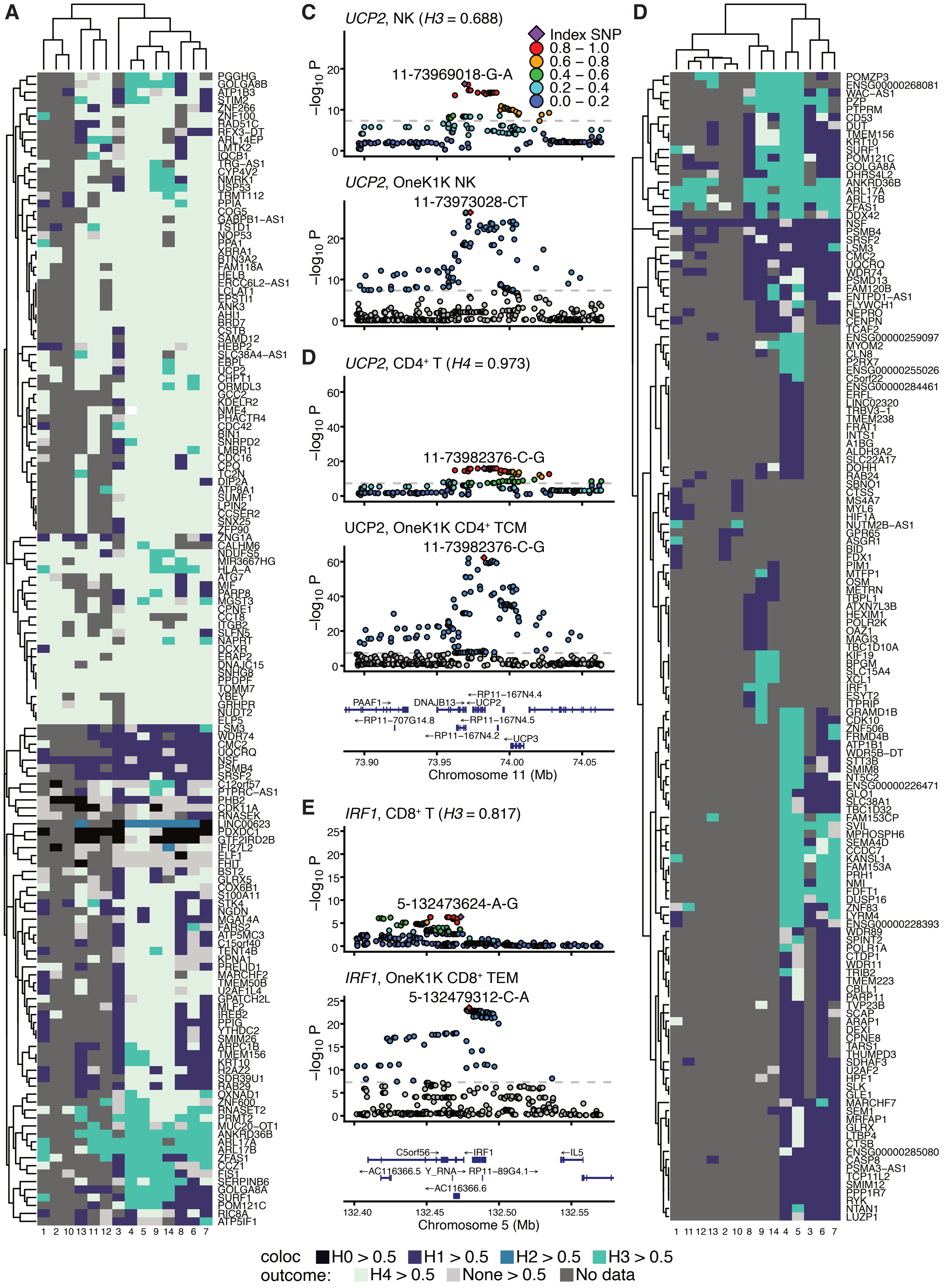
A
C
UCP2, NK (H3 = 0.688)
Index SNP
0.8 – 1.0
0.6 – 0.8
0.4 – 0.6
0.2 – 0.4
0.0 – 0.2
11-73969018-G-A
UCP2, OneK1K NK
11-73973028-CT
D
UCP2, CD4⁺ T (H4 = 0.973)
11-73982376-C-G
UCP2, OneK1K CD4⁺ TCM
11-73982376-C-G
PAAF1→
DNAJB13→
←RP11-167N4.4
←UCP2
←RP11-707G14.8
←RP11-167N4.5
←RP11-167N4.2
←UCP3
73.90
73.95
74.00
74.05
Chromosome 11 (Mb)
E
IRF1, CD8⁺ T (H3 = 0.817)
5-132473624-A-G
IRF1, OneK1K CD8⁺ TEM
5-132479312-C-A
C5orf56→
←IRF1
←IL5
←AC116366.5
Y_RNA→
RP11-89G4.1→
←AC116366.6
132.40
132.45
132.50
132.55
Chromosome 5 (Mb)
D
coloc outcome:
H0 > 0.5
H1 > 0.5
H2 > 0.5
H3 > 0.5
H4 > 0.5
None > 0.5
No data

**Supplementary Figure 3 *(previous page)*.** Colocalisation results between Indonesia and OneK1K. **A)** Summary of colocalisation results for all eGenes tested for colocalisation at least 9 times. Column labels refer to the numeric labels in Figure 3C. **B)** Summary of colocalisation results for eGenes where shared causality between Indonesia and OneK1K1 was not supported across multiple cell types. All eGenes tested for colocalisation at least 2 times and where shared causality was supported under 25% of the time are shown. Column labels refer to the numeric labels in **C**. **C)**-**E)** Examples of colocalisation against OneK1K data. Indonesian data is shown on the top and OneK1K on the bottom for **C)** *IRF1* in Indonesian CD8+ T cells and OneK1K CD8+ TEM cells; **D)** *UCP2* in Indonesian $CD4^{+}$ T and OneK1K CD4+ TCM cells and **E)** *UCP2* in NK cells. Pairwise LD is not available for OneK1K; instead, red indicates the index SNP and blue points exceed nominal genome-wide significance.

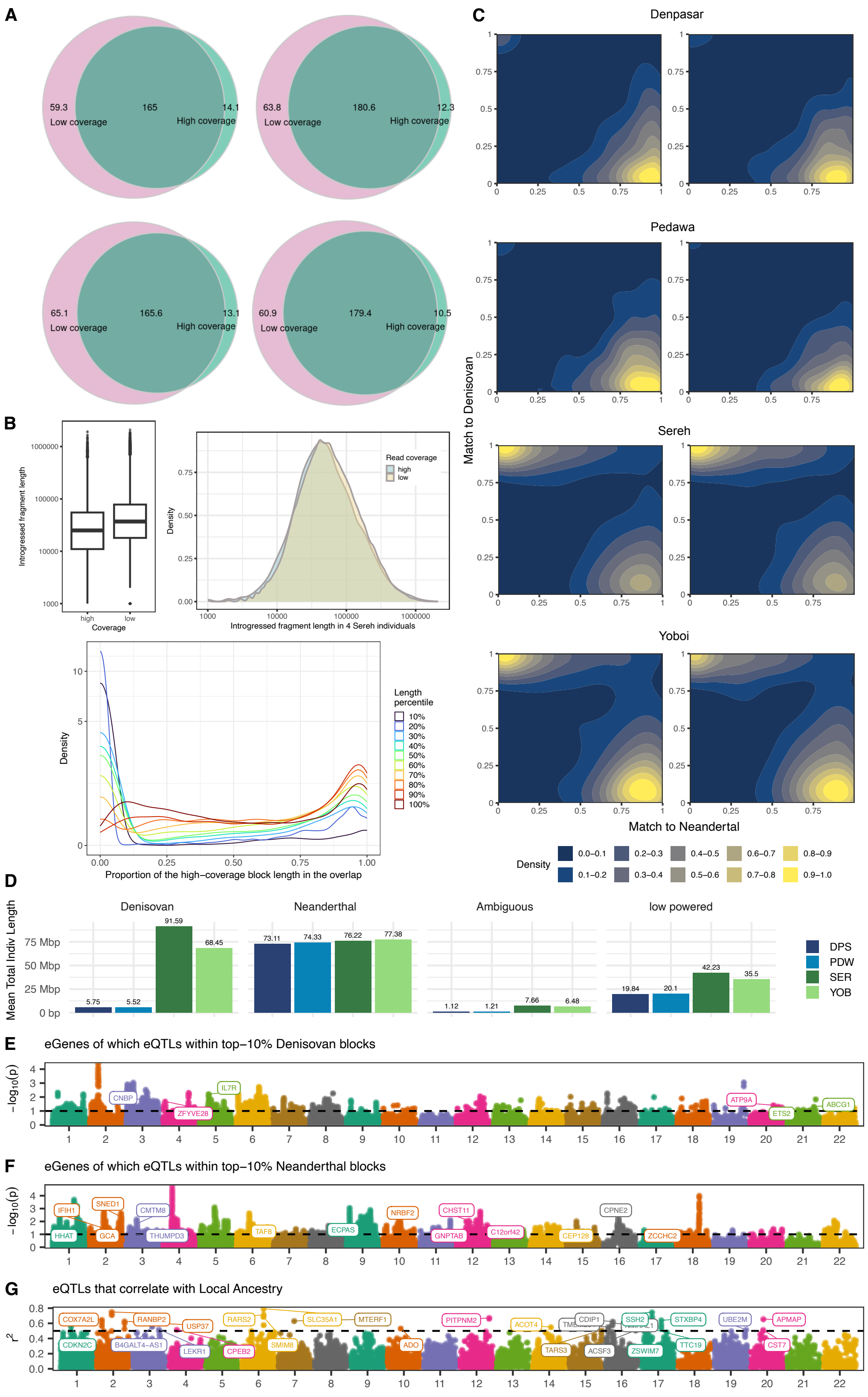
A
59.3
Low coverage
165
14.1
High coverage
63.8
180.6
12.3
65.1
165.6
13.1
60.9
179.4
10.5
B
Introgressed fragment length
Coverage
high
low
Density
Read coverage
Introgressed fragment length in 4 Sereh individuals
Length percentile
10%
20%
30%
40%
50%
60%
70%
80%
90%
100%
Proportion of the high−coverage block length in the overlap
C
Denpasar
Pedawa
Sereh
Yoboi
Match to Denisovan
Match to Neandertal
Density
0.0−0.1
0.1−0.2
0.2−0.3
0.3−0.4
0.4−0.5
0.5−0.6
0.6−0.7
0.7−0.8
0.8−0.9
0.9−1.0
D
Denisovan
Neanderthal
Ambiguous
low powered
Mean Total Indiv Length
75 Mbp
50 Mbp
25 Mbp
0 bp
5.75
5.52
91.59
68.45
73.11
74.33
76.22
77.38
1.12
1.21
7.66
6.48
19.84
20.1
42.23
35.5
DPS
PDW
SER
YOB
E
eGenes of which eQTLs within top−10% Denisovan blocks
CNBP
IL7R
ZFYVE28
ATP9A
ETS2
ABCG1
F
eGenes of which eQTLs within top−10% Neanderthal blocks
IFIH1
SNED1
CMTM8
HHAT
GCA
THUMPD3
TAF8
ECPAS
NRBF2
CHST11
GNPTAB
C12orf42
CEP128
CPNE2
ZCCHC2
G
eQTLs that correlate with Local Ancestry
COX7A2L
RANBP2
USP37
RARS2
SLC35A1
MTERF1
PITPNM2
ACOT4
CDIP1
SSH2
STXBP4
UBE2M
APMAP
CDKN2C
B4GALT4−AS1
LEKR1
CPEB2
SMIM8
ADO
TARS3
ACSF3
ZSWIM7
TTC19
CST7

**Supplementary Figure 4 *(previous page)*.** Archaic introgression and local ancestry inference in low-coverage whole genome sequencing data **A)** Overlap (in Mbp) between introgressed segments inferred in the same 4 individuals from Sereh sequenced at low and high coverage. For the ease of visualisation, here introgression was inferred with unphased option in HMMIX (identifies introgression common between two haplotypes in each individual). **B)** Introgressed segment length distribution in high and low coverage data for 4 Sereh individuals. Longer segments tend to have bigger overlap with segments inferred in high coverage data (one representative individual shown). **C)** Contour plots showing normalised density of match proportion to Neanderthal (average of Vindija, Chagyrskaya, Altai Neanderthals) and Denisovan genomes in the introgressed segments. Two representative individuals from each population are shown. **D)** The amount of inferred introgression in each population. **E)** Manhattan plot of Denisovan-introgressed segments in the dataset ranked by frequency, with eGenes within the top-10% introgressed segments are highlighted. **F)** Manhattan plot of Neanderthal-introgressed segments in the dataset ranked by frequency, with eGenes within the top-10% introgressed segments are highlighted. **G)** Manhattan-like plot of eSNPs that are associated with local ancestry ($r^2$ > 0.5).

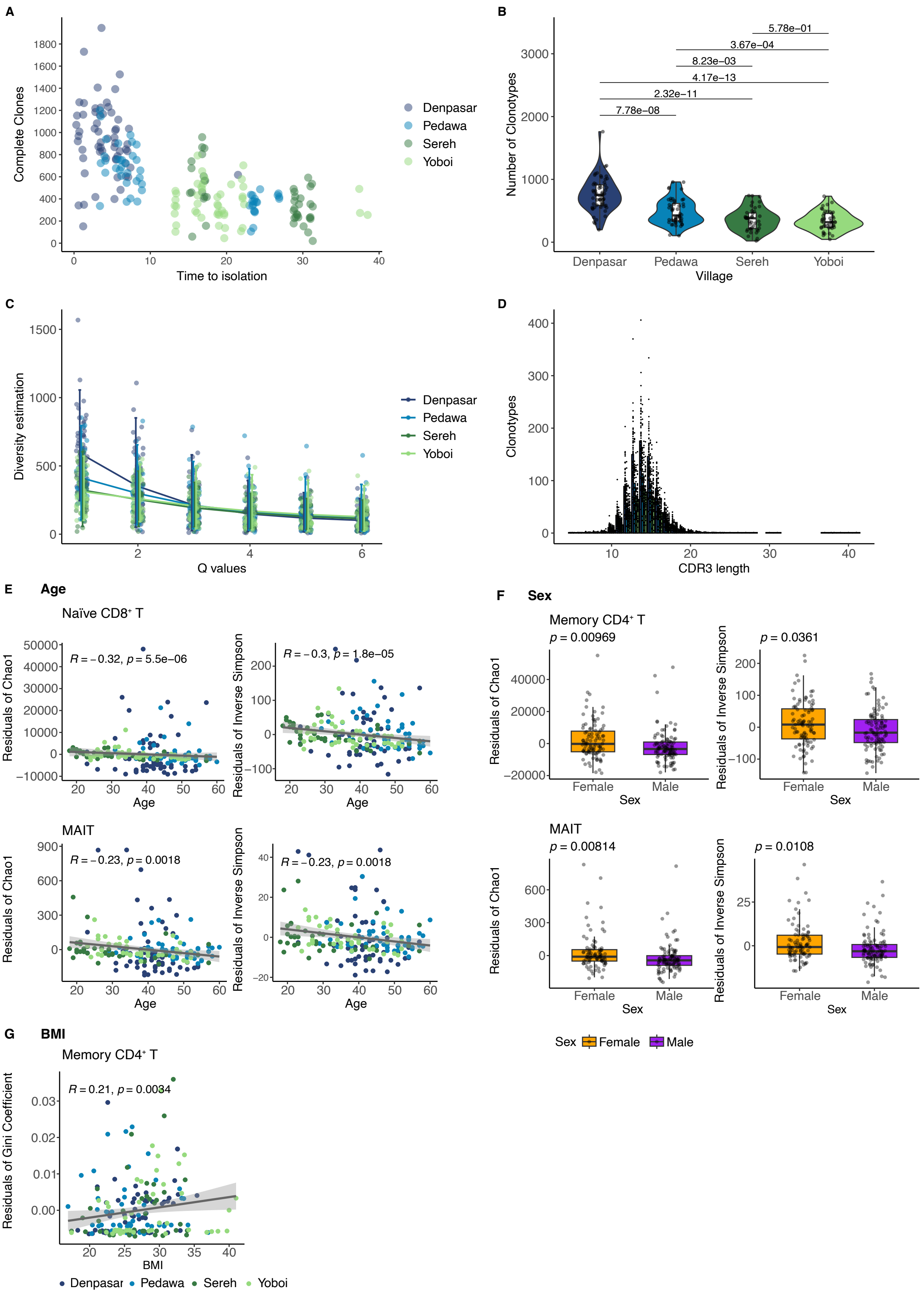
A
Complete Clones
Time to isolation
Denpasar
Pedawa
Sereh
Yoboi
B
Number of Clonotypes
Village
5.78e−01
3.67e−04
8.23e−03
4.17e−13
2.32e−11
7.78e−08
C
Diversity estimation
Q values
D
Clonotypes
CDR3 length
E Age
Naïve CD8⁺ T
Residuals of Chao1
Residuals of Inverse Simpson
R = −0.32, p = 5.5e−06
R = −0.3, p = 1.8e−05
MAIT
R = −0.23, p = 0.0018
R = −0.23, p = 0.0018
Age
F Sex
Memory CD4⁺ T
p = 0.00969
p = 0.0361
MAIT
p = 0.00814
p = 0.0108
Female
Male
Sex
G BMI
Memory CD4⁺ T
Residuals of Gini Coefficient
R = 0.21, p = 0.0034
BMI

**Supplementary Figure 5** ***(previous page)*****.** Single cell TCR repertoire data exploration within each village **A)** Correlation between number of cells with complete TCR repertoire and time to isolation. **B)** Pairwise comparisons between villages were tested using Wilcoxon-rank sum test and adjust multiple comparisons using Benjamin-Hochberg. **C)** Hill diversity across all donors to observe number of clones and clonal distribution. **D)** Distribution of CDR3 length for each donors across all cell types. **E-G)** TCR diversity was estimated by Chao1, Inverse Simpson, and Gini coefficient. Associations between diversity and metadata, including **E)** age, **F)** sex, and **G)** BMI, were examined. Naïve $CD8^+$ T cells and MAIT cells repertoires showed debrease diversity with donor age. To control confounding effects, diversity indices were fitted in linear models regressing out sex, age, and time to isolation. Only significant associations ($p < 0.05$) are shown.

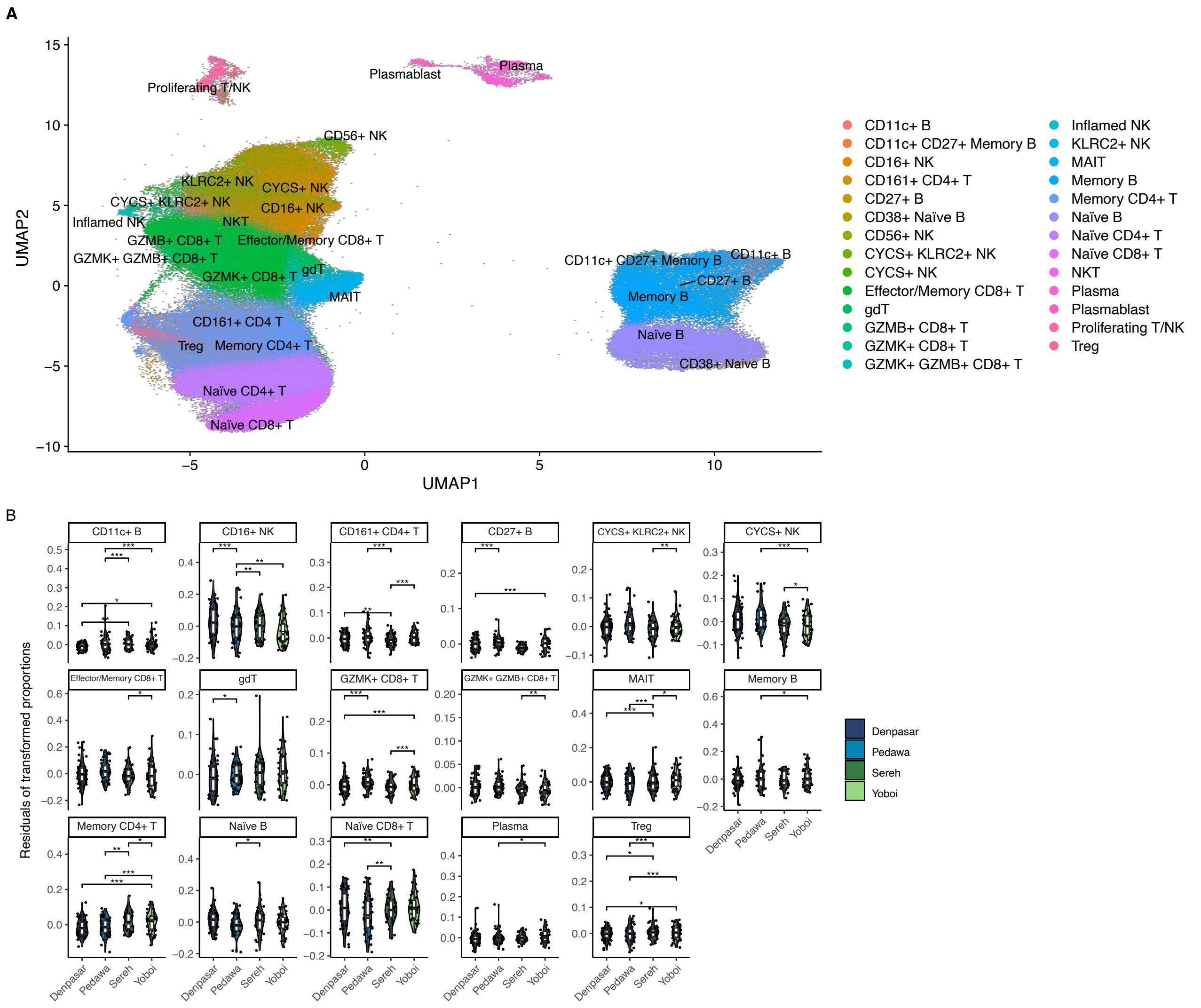


**Supplementary Figure 6.** Deeper cell types annotation and residuals of proportion in lymphoid cells **A)** UMAP of lymphoid lineages colored by level 3 **B)** Violin plots of cell type proportions after adjusting by age, sex, and time to isolation. Only cell types showing significant differences are displayed. Statistical tests were performed using one-way ANOVA, adjusted by multiple testing using Benjamin-Hochberg.

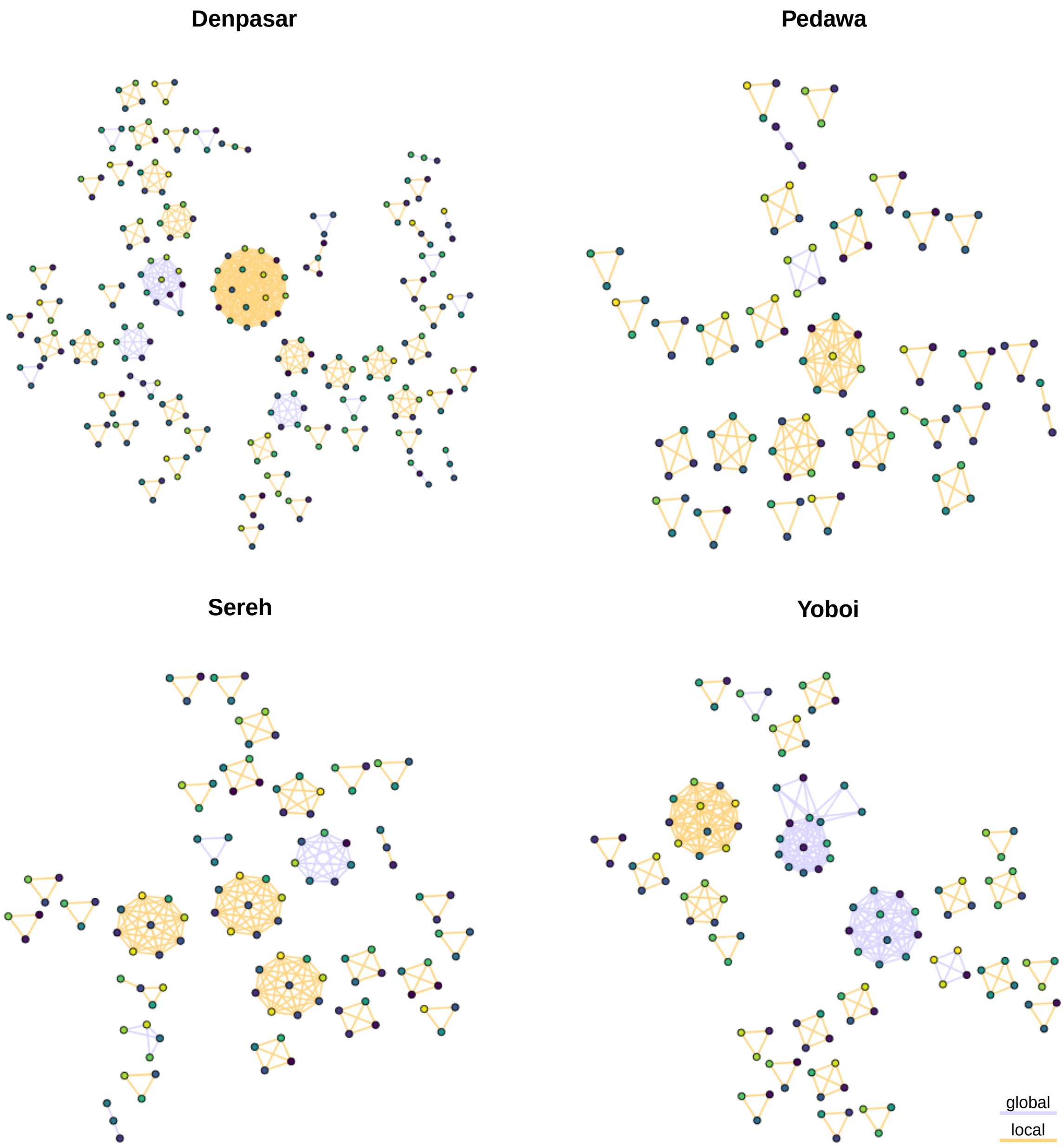


**Supplementary Figure 7.** Motif enrichment within each village in CD8⁺ T cells. Networks of global (purple) and local (orange) clusters analysed by GLIPH2 based on CDR3 sequences of TCR$\beta$ chain. Each node represents a clonotype with colours indicating donors. Displayed clusters correspond to those that filtered Fisher's exact test for significance ($p < 0.05$).

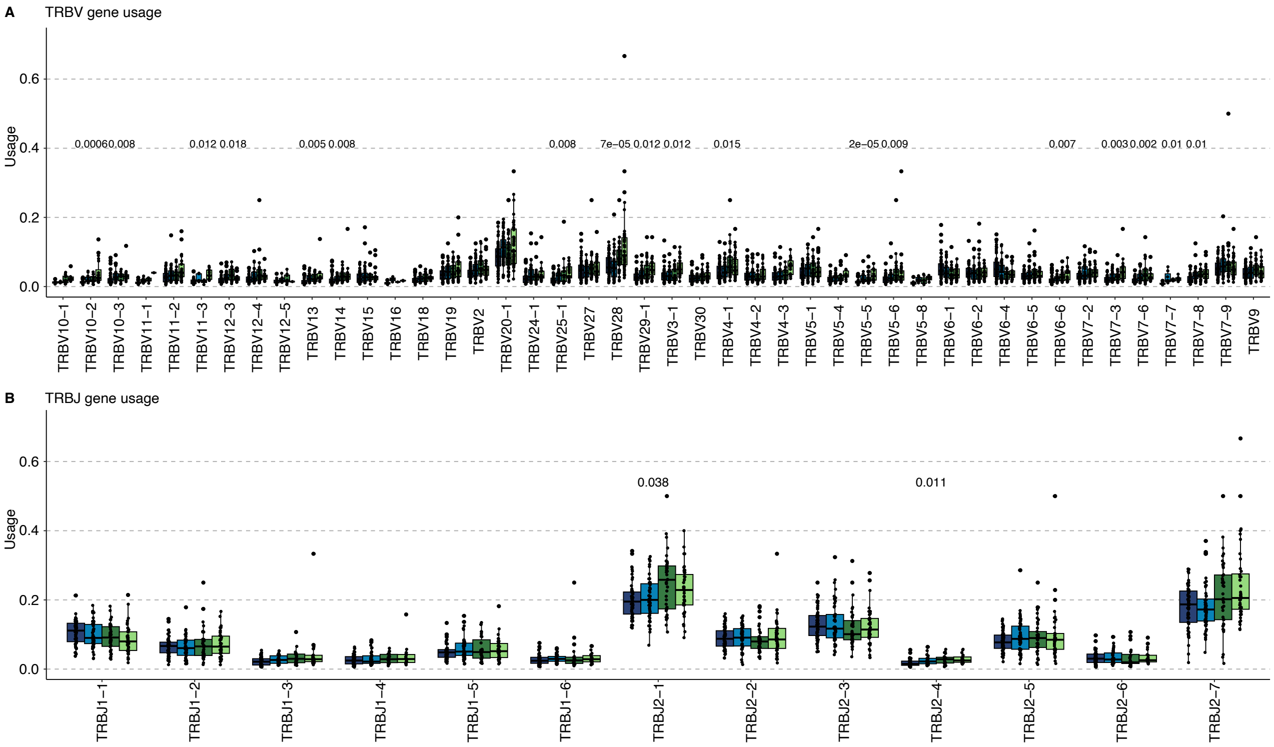


**Supplementary Figure 8.** TRBV and TRBJ gene usage in CD8$^{+}$ cells Box plots depict distribution of **A)** TRBV and **B)** TRBJ gene segment usage individually. Statistical tests were performed using FDR-corrected Kruskal-Wallis $p < 0.05$

### Supplementary Tables

**Table S1.** Cellranger QC summary (per batch).

**Table S2.** Per-donor information.

**Table S3.** Marker genes for cell type annotation.

**Table S4.** Estimated pairwise kinship between all individuals in the study.

**Table S5.** Number of eQTLs and eGenes identified after quasar and mashr.

**Table S6.** Colocalisation results between Indonesia and OneK1K for all tested eQTLs.

**Table S7.** Genes with significant GxE interactions.

**Table S8.** eQTLs related to local ancestry.

**Table S9.** eQTLs related to archaics introgression.

**Table S10.** NetRep module preservation p-values.

**Table S11.** GO Biological Process set enrichment analysis for each module.

**Table S12.** Hallmark gene set enrichment analysis for each module.

**Table S13.** DE pairwise result by five cell types

**Table S14.** Viral-related GO terms in expanded and unexpanded $CD8^+$ T cells

**Table S15.** Epitope annotation in significant global motifs.
